# Mindfulness meditators show enhanced working memory performance concurrent with different brain region engagement patterns during recall

**DOI:** 10.1101/801746

**Authors:** NW Bailey, G Freedman, K Raj, KN Spierings, LR Piccoli, CM Sullivan, SW Chung, AT Hill, NC Rogasch, PB Fitzgerald

**Affiliations:** Epworth Centre for Innovation in Mental Health, Epworth Healthcare, The Epworth Clinic, Camberwell, Victoria, Australia, 3004; Monash Alfred Psychiatry Research Centre, Monash University Central Clinical School, Commercial Rd, Melbourne, Victoria, Australia; Brain, Mind and Society Research Hub, School of Psychological Sciences, Monash Biomedical Imaging, Turner Institute for Brain and Mental Health, Monash University, VIC, Australia; Temerty Centre for Therapeutic Brain Intervention, Centre for Addiction and Mental Health, University of Toronto, Toronto, Ontario, Canada; Discipline of Psychiatry, Adelaide Medical School, University of Adelaide, Adelaide, SA, Australia; South Australian Health and Medical Research Institute (SAHMRI), Adelaide, SA, Australia

**Keywords:** Attention, Mindfulness Meditation, EEG, Working Memory, Alpha, Theta, P3, FN400, parietal old/new effect, 1/f aperiodic activity

## Abstract

Mindfulness meditation has been shown to improve working memory (WM). However, the altered brain activity underpinning these improvements is underexplored. In non-meditating individuals, modulation of theta and alpha oscillations and 1/f aperiodic activity during WM has been found to be related to WM performance. Resting theta and alpha oscillations have been found to differ in meditators, but WM related oscillation changes and 1/f aperiodic activity have not yet been examined. Additionally, WM event-related-potentials (ERPs) are modulated by attention, which is also enhanced by meditation, so these neural measures are candidates for exploring neural activity underpinning WM improvement in meditators. We recorded EEG from 29 controls and 29 meditators during a modified Sternberg WM task and compared theta, alpha, and 1/f aperiodic activity during the WM delay, and ERPs time-locked to the WM probe. Meditators responded more accurately (p = 0.008, Cohen’s d = 0.688). Meditators also showed different ERP distributions with earlier left-temporal activation and more frontal distribution of activity (FDR-p = 0.0186, η^2^ = 0.0903), as well as a reduction in overall neural response strength (FDR-p = 0.0098, η^2^ = 0.1251). While a higher proportion of meditators showed theta oscillations during the WM delay, no other differences in theta, alpha or 1/f aperiodic activity were present. These results suggest that increased WM performance in meditators might not be the result of higher amplitudes of typical WM activity, but instead due to an alternative neural strategy during WM decision making, which may allow more accurate responses with less neural activation.

**Highlights:**

- Long term mindfulness meditators showed improved working memory (WM) accuracy
- This was concurrent with earlier left temporal activation following probe stimuli
- As well as a more frontal distribution and reduced overall neural response strength
- No oscillation differences were present in the working memory delay period
- Improved WM from altered neural strategy rather than increased neural activity

## Introduction

Mindfulness meditation has been shown to improve attention and alter neural activity related to attention (Lutz, Slagter, Dunne, & Davidson, 2008; MacLean et al., 2010; Tang et al., 2007). Attentional processes are essential for working memory (WM) performance, which is a capacity limited system that enables behavioural responses in the present to be informed by stimuli presented recently but currently not available for sensory processing (Baddeley, 2012; Christophel, Klink, Spitzer, Roelfsema, & Haynes, 2017; Ma, Husain, & Bays, 2014; Wolpaw, 2002). In addition to the effect of meditation on attention, a smaller amount of research has shown that meditation has a positive impact on WM (Jha, Stanley, Kiyonaga, Wong, & Gelfand, 2010; Mrazek, Franklin, Phillips, Baird, & Schooler, 2013; Quach, Mano, & Alexander, 2016; Van Vugt & Jha, 2011; Zeidan, Johnson, Diamond, David, & Goolkasian, 2010). However, very little research has examined neural activity underpinning this improved WM performance in meditators. There are a number of reasons to be interested in exploring the neural changes responsible for these meditation-related WM improvements. Firstly, better understanding of WM related neural activity in meditators could be informative regarding whether meditation simply leads to an enhancement in attention, and the attention enhancement is solely responsible for all other cognitive enhancements, or whether there are domain specific enhancements as well (Buttle, 2011; Jha et al., 2010) (see supplementary materials for a detailed discussion of this point). Expanding from this point, the research may be informative of how meditation works at a neural level, and as a result, WM research in meditation might lead to a biomarker to measure improvements as a result of meditation. This might enable improved interventions that more directly target the mechanism of action, and potential predictors of which clinical groups or individuals might benefit most from mindfulness meditation (Britton et al., 2018). Thirdly, given that meditators do show enhanced WM performance, understanding neural activity underpinning WM performance in meditators could be informative regarding neural activity underlying good WM function.

In order to explore the effects of meditation on WM, neural activity related to WM can be examined using a modified Sternberg task (Sternberg, 1966), which presents a WM set (referred to henceforth as the ‘WM set presentation period’, which in the case of the current study consisted of eight simultaneously presented visual letters), followed by a ‘WM delay period’ (which was a blank screen in current study), followed by a WM probe period (a single probe letter, which may or may not have been in the memory set, see supplementary materials for a detailed explanation of the selection of WM period labels). To complete the modified Sternberg task, participants are instructed to push one button if the probe letter was in the WM set, and another button if it was not. Separating the different periods of WM in this way allows for separate analysis of neural processes related to specific WM relevant functions, useful for fully characterising the effects of mindfulness on WM.

In research on non-meditators, brain regions associated with WM processing are prefrontal, parietal, and medial temporal regions, with other regions recruited for sensory modality specific WM functions (Christophel et al., 2017; Gu, van Rijn, & Meck, 2015). Oscillatory activity is modulated in these brain regions during performance of WM functions. In particular, neural activity measured with EEG during the WM delay period of the Sternberg task shows increased theta activity (4-8 Hz) in fronto-midline electrodes (Brookes et al., 2011; Kottlow et al., 2015; L. Payne & Kounios, 2009). This activity has been suggested to be generated by the anterior cingulate cortex (ACC), and is thought to reflect an attentional mechanism enabling selection of specific task relevant neural processes for enhancement from multiple competing neural processes (Cavanagh & Frank, 2014; Sauseng, Griesmayr, Freunberger, & Klimesch, 2010; Sauseng, Hoppe, Klimesch, Gerloff, & Hummel, 2007). Increased theta power has been associated with increased performance both between individuals (Maurer et al., 2015) and within individuals across trials (Scheeringa et al., 2009) as well as being associated with increased WM loads (O. Jensen & Tesche, 2002). Previous research has also shown that theta activity modulations are associated with meditation (Tang et al., 2009)

In addition to selecting for neural processes involved in WM functioning, non-relevant regions need to be inhibited to enable WM performance. Alpha oscillations have been suggested by previous research to reflect these inhibitory processes, and in the Sternberg task neural activity during the WM delay period shows increased upper alpha power (10-12.5 Hz) in parieto-occipital electrodes (Wolfgang Klimesch, Sauseng, & Hanslmayr, 2007). This sustained high power alpha activity has been suggested to reflect suppression of cortical gain (in contrast to low power bursting alpha which may enhance gain) (Peterson & Voytek, 2017). As such, high power alpha may reflect top-down inhibition of visual processing brain regions that are not relevant processes required in the WM delay period, as visual information processing is not required during the WM delay period (a blank screen) (Wolfgang Klimesch et al., 2007). Increased alpha activity in these regions is associated with decreased fMRI blood flow measured in the visual cortex, reflecting the suppression of potentially disruptive activity in this region (Scheeringa et al., 2009). This suppression of non-relevant brain regions leads to facilitation of WM performance. On a single trial basis, trials with higher alpha power are more likely to be followed by a correct response (Scheeringa et al., 2009) and larger WM set sizes (higher WM loads with more stimuli to remember) are associated with increased alpha power (O. Jensen & Tesche, 2002). Previous research has also indicated that meditators are better able to modulate alpha activity in brain regions processing (tactile) distractors (Kerr, Sacchet, Lazar, Moore, & Jones, 2013; Lagopoulos et al., 2009; Tang et al., 2009). These findings suggest that an increase in the ability to modulate alpha and theta oscillations during WM delay periods might reflect an increase in attentional mechanisms underpinning task adaptation, leading to increased performance in meditators.

In addition to examining oscillatory activity, recent research has shown that while traditional measures of oscillatory power include aperiodic (non-oscillatory) neural activity, it is important to separate oscillatory power from aperiodic neural activity (which shows a ‘1 divided by the frequency value’ distribution, termed ‘1/f aperiodic activity’) when assessing the functional relevance of neural oscillations (Haller et al., 2018). Without this separation, significant differences in measures of power within a specific oscillation frequency may reflect differences in the 1/f aperiodic activity rather than a difference in the oscillation (Haller et al., 2018). This research has also shown that the slope of the 1/f aperiodic activity reflects neural activity that is not rhythmic or oscillatory but is suggested to be produced by the Poisson distribution of spiking synaptic potential timing (where the average rate of synaptic potentials is constant, but the time between potentials is variable and of similar amplitude to the mean), with steeper 1/f aperiodic activity slopes perhaps reflecting reduced excitation / inhibition ratios (driven by glutamate and GABA respectively) (Gao, Peterson, & Voytek, 2017). The excitation / inhibition balance has been suggested to be vital for information transmission and gating (Gao et al., 2017). Shallower 1/f aperiodic activity slopes are related to WM performance decline in aging while steeper slopes are better than oscillatory power as a predictor of schizophrenia (Gao et al., 2017; Peterson, Rosen, Campbell, Belger, & Voytek, 2018; Voytek et al., 2015). The 1/f aperiodic activity offset has also been shown to be related to overall neuronal firing rates / spiking activity (Manning, Jacobs, Fried, & Kahana, 2009; Miller et al., 2012) and the fMRI bold signal (Winawer et al., 2013). As such, measures of 1/f aperiodic activity are likely to be of interest when examining WM related neural activity in meditators.

In addition to neural activity during WM delay periods, a significant amount of research has examined neural activity associated with memory probe presentation. This research has typically focused on neural activity time-locked to the onset of the probe and averaged over many presentations of probe stimuli. This approach eliminates neural activity that is not consistently time locked to the WM probe, thus comparing neural activity only associated with probe stimulus processing, assumed to reflect activity associated with WM decisions (Friedman & Johnson Jr, 2000). The measurement of neural activity resulting from this time-locking to a stimulus and averaging is referred to as an event-related potential (ERP). Previous memory research has shown memory probe related ERPs with positive voltages in central/lateral parietal regions, surrounded by negative voltages (maximal in fronto-central regions) from 300 to 700 ms following stimuli presentation. The ERP has been given different names by different researchers and has been subdivided in different ways depending on task demands and time periods of analysis.

Neural activity earlier in the WM probe period ERP window (from 300 to 450 ms following stimuli presentation) has been referred to as the FN400 (FN referring to ‘frontal negativity’ due to the negative voltages in frontal regions that are typically compared, although it is worth noting that positive activity is typically present in parietal regions) (Curran & Cleary, 2003). The FN400 has been associated with familiarity (Curran & Cleary, 2003; Duarte, Ranganath, Winward, Hayward, & Knight, 2004), conceptual processing (Woodruff, Hayama, & Rugg, 2006) and semantic processing (Federmeier & Laszlo, 2009; Yonelinas, 2002). It is thought to be generated in part by left temporal regions, and FN400 activity in those regions are related to recognition (Stróżak, Abedzadeh, & Curran, 2016). The FN400 is also modulated by instructions about which parts of stimuli to attend to, suggesting it is related to attentional processes (Rugg & Curran, 2007). Research has shown that larger FN400 amplitudes are associated with increased familiarity and memory strength (Finnigan, Humphreys, Dennis, & Geffen, 2002; Rugg et al., 1998). Neural activity later in the WM probe period ERP (450 to 700 ms) typically shows more positive voltages in central and lateral parietal regions when participants have previously seen stimuli than when they have not seen the stimuli before, so the ERP has been referred to as the parietal old/new effect (Stróżak et al., 2016). This activity has been suggested to be associated with conscious recollection (Duarte et al., 2004; Rugg & Curran, 2007; Woodruff et al., 2006; Yonelinas, 2002). The amplitude of the parietal old/new effect has been suggested to reflect attention orientation to recollection (Rugg & Henson, 2002; Wagner, Shannon, Kahn, & Buckner, 2005) and is associated with accuracy, or confidence in the accuracy of the response (Finnigan et al., 2002). Research has suggested that the earlier processes reflected by the FN400 are less effortful than the later processes reflected by the parietal old/new effect (Rugg & Curran, 2007).

When WM ERPs are measured specifically in the Sternberg WM task, activity across this WM probe period ERP window has been referred to as the P3 (which shows a similar time window and distribution to both the FN400 and the parietal old/new effect) (Chang, Huang, Chen, & Hung, 2013; Ergen, Yildirim, Uslu, Gürvit, & Demiralp, 2012). The WM P3 has been suggested to reflect a process that inhibits widespread cortical regions, suppressing irrelevant neural activity from interrupting the WM relevant processes (W Klimesch, Doppelmayr, Schwaiger, Winkler, & Gruber, 2000). In general, P3 amplitudes are thought to indicate attentional resource allocation to stimulus processing while P3 latency is thought to reflect processing speed (Pontifex, Hillman, & Polich, 2009). Researchers have suggested the P3 indexes WM retrieval, and that it is related to memory scanning and decision making (Ergen et al., 2012). Lastly, it is worth noting that most of the WM ERP research has measured activity from single or small clusters of electrodes. In contrast to this approach, measuring the scalp distribution using all electrodes is informative regarding the engagement of different brain regions, indicating different functional engagement underlying cognition (Friedman & Johnson Jr, 2000). Research by our lab using other cognitive tasks has shown that mindfulness meditators showed altered distributions of neural activity, typically with more frontal distributions (N. W. Bailey et al., 2019; Wang et al., 2019). Our research has also demonstrated reduced overall neural response strength concurrent with increased behavioural performance in meditators (N. W. Bailey et al., 2019; Wang et al., 2019).

The aim of the current research was to examine both the distribution and overall strength of neural activity related to both WM delay and WM probe periods in long term mindfulness meditators compared to healthy (demographically matched) non-meditators. It was hypothesised a priori that 1) meditators would show increased parieto-occipital alpha and fronto-midline theta power during the WM delay period, reflecting increased attentional modulation of top-down inhibitory and executive control functions, 2) meditators would show reduced ERP activity when measured across all electrodes, and 3) meditators would show a more frontal distribution of the WM ERPs. Exploratory post-hoc comparisons were made of the 1/f aperiodic activity slope and offset parameters, as well as of alpha and theta oscillations separated from the 1/f aperiodic activity. Lastly, in replication of previous research, we had a confirmation hypothesis that the meditation group would show higher WM accuracy than the control group.

## Methods

### Participants and Self-Report Data

Thirty-four meditators and 36 demographically matched non-meditating controls were recruited through community advertising. Inclusion criteria for meditators consisted of having a current meditation practice involving at least two hours per week of practice, with at least six months of meditation experience (all meditators except three had more than two years of meditation experience). Phone screening and in-person interviews were administered by experienced mindfulness researchers (GF, KR, NWB) to ensure meditation practices were mindfulness-based, using Kabat-Zinn’s definition “paying attention in a particular way: on purpose, in the present moment, and nonjudgmentally” (Kabat-Zinn, 2009). Further screening ensured meditation practices were consistent with either focused attention on the breath or body-scan. Uncertainties were resolved by consensus between two researchers including the principal researcher (NWB). Control group participants were screened to ensure they had less than two hours of lifetime experience with any kind of meditation. Exclusion criteria involved self-reported current or historical mental or neurological illness, or current psychoactive medication or recreational drug use. Participants were interviewed with the Mini International Neuropsychiatric Interview for DSM-IV (Sheehan et al., 1998) and excluded if they met criteria for any DSM-IV illness. Participants were also excluded if they scored in the mild or above range in the Beck Anxiety Inventory (BAI) (Steer & Beck, 1997) or Beck Depression Inventory (Beck, Steer, & Brown, 1996). All participants were between 19 and 62 years of age and had normal or corrected to normal vision.

All participants provided informed consent prior to participation. The study was approved by the Ethics Committee of the Alfred Hospital and Monash University (approval number 194/14), and conducted in accordance with the Declaration of Helsinki. At the beginning of the testing session participants completed demographic and self-report forms including their age, gender, years of education, handedness, and estimated how many years they had been practicing meditation for, and the average number of minutes per week they spent meditating in the last two months. Participants completed the Freiburg Mindfulness Inventory (FMI) (Walach, Buchheld, Buttenmüller, Kleinknecht, & Schmidt, 2006), Five Facet Mindfulness Questionnaire (FFMQ) (Baer, Smith, Hopkins, Krietemeyer, & Toney, 2006), BAI and BDI-II (see Table 1 for a summary of the self-report data). Prior to the modified Sternberg WM task, participants completed a Go/Nogo task (N. W. Bailey et al., 2019), colour Stroop task, emotional Stroop task (Marcu et al. in preparation), and Nback task (Wang et al., 2019), which took approximately one hour to complete in total including breaks within and between tasks.

**Table 1.** Means, standard deviations and statistics for demographic and self-report comparisons between groups.

|  | Controls<br>M (SD) | Meditators<br>M (SD) | Statistics |
| --- | --- | --- | --- |
| Age | 36.45 (13.92) | 37.31 (11.50) | $t(56) = 0.257, p = 0.798$ |
| Gender (F/M) | 19/10 | 17/12 | $\chi^2 = 0.293, p = 0.588$ |
| Years of Education | 15.89 (3.22) | 16.90 (2.74) | $t(56) = 1.269, p = 0.210,$ |
| Preferred Hand (R/L) | 28/1 | 27/2 | $\chi^2 = 0.352, p = 0.553$ |
| Years Meditating |  | 9.22 (10.87) |  |
| Hours Meditating/Week |  | 5.41 (4.11) |  |
| BAI | 4.62 (5.45) | 3.62 (4.30) | $t(56) = 0.775, p = 0.442$ |
| BDI-II | 1.69 (2.62) | 1.24 (2.21) | $t(56) = 0.704, p = 0.485$ |
| FMI | 39.90 (8.38) | 45.18 (6.87) | $t(56) = 2.597, p = 0.012$ |
| FFMQ | 137.69 (12.67) | 152.41 (17.60) | $t(56) = 3.656, p = 0.001$ |

Data from four control participants were excluded after scoring in the mild depression range on the BDI-II. Data from three control participants were excluded from EEG analysis due to poor EEG signal quality. WM EEG data were not collected from five meditators due to time constraints. Accuracy data from two control and three meditator participants were excluded due to an intermittent button fault during those sessions resulting in unreliable accuracy measurement (while EEG and reaction time data were included as a sufficient number of correctly responded to epochs were still provided). Following exclusions, 29 control and 29 meditator participants were included in the EEG and reaction time analyses, and 27 control and 26 meditator participants were included in accuracy analysis.

### Data Availability

Participants involved in the study did not provide consent to data sharing, and data sharing was not approved by the Alfred Hospital ethics committee, so the results reported in the paper comprise the complete data available for sharing. The Alfred Hospital ethics committee can be contacted via.

### Task and Stimuli

Participants performed a modified Sternberg WM task with eight simultaneously presented letters as stimuli while 64-channel EEG was recorded (see Figure 1) (N. Bailey et al., 2019). Letters were selected from a set of 15 potential consonants (B, C, D, F, H, J, K, L, N, R, S, T, Y, W, and Z), and all stimuli were presented with Neuroscan STIM2 software (Compumedics, Melbourne, Australia). Trials began with a fixation cross (800 ms) followed by a blank screen (1000 ms). The WM stimuli set was then presented (WM set presentation period, 4000 ms) followed by a blank screen for the WM delay period (3000 ms). A single probe letter was then presented (WM probe presentation period, 2000 ms), to which participants had to respond with one button if the probe had been present in the memory set, and another if the probe had not been present in the memory set. There was a 50% probability in each trial that the probe had been present in the memory set. Following the offset of the probe, a brief visual mask was presented (166.67 ms), then a blank screen (1883.33 ms), before the fixation cross was once again presented to begin the next trial. The total task consisted of two blocks with 26 trials per block. Participants performed a brief practice version of the task prior to the recording.

**Figure 1.**
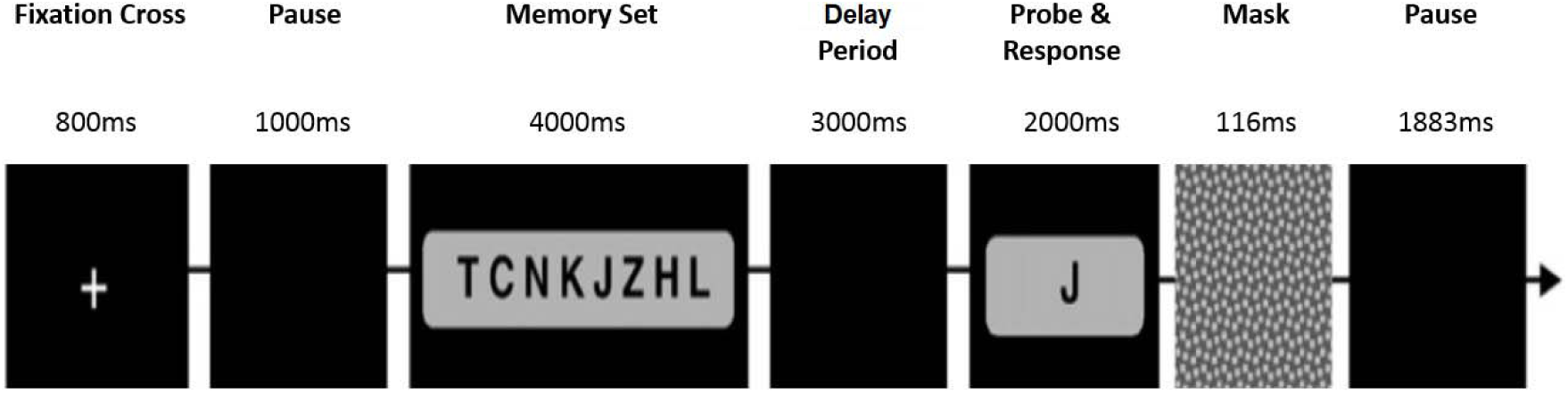
– Task design and stimuli timing for the modified Sternberg task. WM set letters were presented simultaneously, and always contained eight letters. Responses made outside of the WM probe presentation period were considered incorrect.

### Electrophysiological Recording, Pre-Processing, Power and ERP Analysis

A Neuroscan 64-channel Ag/AgCl Quick-Cap was used to acquire EEG data through Neuroscan Acquire software and a SynAmps 2 amplifier (Compumedics, Melbourne, Australia). Electrodes were referenced to an electrode between Cz and CPz. Eye movements were recorded with a supraorbital electrode above the left eye. Electrode impedances were kept below 5kΩ. EEG was recorded at 1000 Hz with an online bandpass filter of 0.05 to 200 Hz. Data were pre-processed offline in MATLAB (The Mathworks, Natick, MA, 2018a) using EEGLAB for pre-processing (sccn.ucsd.edu/eeglab) (Delorme & Makeig, 2004). Data were filtered, epoched to correct trials only, artifacts automatically then manually rejected, submitted to independent component analysis (ICA) and cleaned of eye movements and remaining artifacts, then referenced to an average reference. A minimum of 20 accepted epochs were required for inclusion. Full details of analysis steps are specified in the supplementary materials.

### Baseline Corrected Power and ERP Analysis

In order to compute power measures the EEG data from each accepted epoch for each participant were submitted to a Morlet Wavelet Transform (with 5 oscillation cycles required to derive power estimates at each timepoint) (Tallon-Baudry, Bertrand, Delpuech, & Pernier, 1997). Power was calculated in the upper alpha band (10-12.5 Hz) and theta band (4-8 Hz). Power was baseline corrected to the middle 600 ms of the blank screen that appeared early in the trial (1000 to 1600 ms following the beginning of the trial). Modulation of power was calculated relative to the baseline using the formula: Baseline Corrected Power (BLC Power) = (WM delay period activity – blank screen reference activity) / blank screen reference activity. This formula provides positive values when power is higher in the WM delay period than the blank screen reference period and negative values when power is lower in the WM delay period. Average BLC power was calculated across the entire WM delay period (5800 to 8800 ms after the start of the trial) within both frequency bands, then averaged over trials for each participant. These averaged BLC power values were used to make statistical comparisons between groups. ERP analysis was conducted time-locked to the onset of the probe stimulus. Voltage data from each electrode for −100 to 1000 ms around the probe were baseline corrected to the −100 to −10 ms period, and all epochs from each participant were averaged for ERP analyses. Probe present and probe absent epochs were averaged together to ensure enough epochs were available for analysis. Exploratory analyses were performed on data from probe present and probe absent conditions averaged separately to check that results were not confounded by averaging conditions. These comparisons are reported in the supplementary materials. Source analysis was also conducted to characterise potential generators of the ERP differences between groups without statistical comparison (methodology reported in supplementary materials).

### Statistical Comparisons

Comparisons of self-report and behavioural data were conducted using SPSS version 23 for behavioural and single electrode statistics, with JASP (JASP Team, 2019) to compute eta squared effect size (η^2^) values (with both η^2^ and partial eta squared – η_p_^2^ reported where these values differed) and the Randomised Graphical User Interface (RAGU) for other EEG analyses (Koenig, Kottlow, Stein, & Melie-García, 2011). Independent samples t-tests were used to verify groups did not differ in age, years of education, BAI, BDI-II, and to test whether groups differed in FMI and FFMQ scores. Chi square tests were used to verify groups did not differ in gender or handedness. Because previous research suggested meditators show superior WM performance, a one-sided t-test was used to compare d-prime accuracy between groups. A repeated measures ANOVA was used to compare reaction times across the 2 groups x 2 conditions (probe present or absent). No outliers were present for data tested with traditional statistics, and data tested with traditional statistics met assumptions of normality, sphericity, and homogeneity of variance.

### Primary Comparisons

Primary statistical comparisons between groups in EEG data were performed using RAGU. RAGU uses reference free global field potential (GFP) measures and randomisation statistics to compare neural response strength and scalp field differences across all electrodes and time points without a priori assumptions about locations or time windows showing significant effects. RAGU controls for multiple comparisons in the spatial dimension by collapsing differences to a single scalp difference map value for distribution comparisons and using the GFP for neural response strength comparisons. It controls for multiple comparisons in the time dimension by ranking the length of significant differences in the real data against the randomised data and ensuring the significant periods in the real data are longer than 95% of the randomised data (global duration control), or that the count of significant timepoints in the real data exceeds the count of significant timepoints in 95% of the randomised data (global count control). Further details about RAGU can be found in the supplementary materials and in Koenig et al. (2011). Differences between groups in overall neural response strength (across all electrodes) were compared using the GFP test. Differences between groups in the distribution of activity across the scalp were compared independently of amplitude using the topographical analysis of variance (TANOVA). GFP and TANOVA tests were used to conduct t-test design comparisons comparing ERPs time-locked to the onset of the probe stimuli from −100 to 1000 ms. Post-hoc GFP and TANOVA tests explored significant time periods averaged across time periods of significant effects that passed multiple comparison controls in the time domain.

To make separate between group comparisons of alpha and theta BLC power during the WM delay period, we used t-test designs in RAGU with Root Mean Square (RMS) and TANOVA tests (to separately compare overall neural response strength and distribution of neural activity respectively). It should be noted that when frequency comparisons are computed with RAGU, the average reference is not computed on the frequency transformed data (the average reference was computed prior to the frequency transforms). As such, the test is a comparison of the RMS between groups, a measure which is a valid indicator of neural response strength in the frequency domain. In other respects, the statistic used to compare RMS between groups is identical to the GFP test (T. Koenig 2018, Department of Psychiatric Neurophysiology, University Hospital of Psychiatry, personal communication). η^2^ effect sizes were computed in RAGU for all comparisons of interest. In order to control for multiple comparisons across all primary hypotheses the Benjamini and Hochberg false discovery rate (FDR) was applied to the global count p-value for each primary statistical comparison (WM delay period theta and alpha RMS and TANOVA, and WM probe period ERP GFP and TANOVA comparisons) (Benjamini & Hochberg, 1995). It was also separately applied to p-values obtained for the average difference between groups during time periods of significance to control for multiple comparisons when focusing on time windows of interest. To enable comparison with other research, both corrected and uncorrected p-values are reported for significant comparisons (labelled ‘FDR-p’ and ‘p’ respectively).

### Exploratory Analyses

Exploratory analyses were not corrected for multiple comparisons, and as such should be taken as preliminary. To assess a potential relationship between neural activity and behaviour, significant periods from group GFP and TANOVA comparisons were averaged and compared to d-prime scores using the GFP covariance and TANCOVA tests. Methods for separating 1/f aperiodic activity and oscillatory activity are described in the supplementary materials and in Haller et al. (2018). T-test comparisons were conducted using SPSS for the following measures at Fz: theta (4 to 8 Hz) centre frequency, bandwidth, amplitude, 1/f aperiodic slope, 1/f aperiodic offset. Repeated measures ANOVA comparisons were conducted including PO7 and PO8 for alpha (8 to 13 Hz) centre frequency, bandwidth, amplitude, 1/f aperiodic slope and 1/f aperiodic offset. Chi-squared tests were also used to determine whether groups differed in the number of participants showing presence/absence of theta and alpha peaks at electrodes of interest (since not all participants showed peaks in each band once the 1/f aperiodic activity was removed). Lastly, microstates were also used to explore significant results obtained in the TANOVA analysis and to justify selections of windows of analysis for TANCOVA and source analyses. Microstate analyses are reported in the supplementary materials.

## Results

### Demographic and Behavioural Comparisons

No differences between groups were present for age, gender, years of education, BAI scores, BDI-II scores, and preferred hand (all p > 0.10). Significant differences were present for FMI and FFMQ scores (p = 0.012 and p = 0.001 respectively). Table 1 reports means, standard deviations and statistics for these comparisons in detail.

D-prime comparisons showed a significant difference, with meditators showing higher performance (t(51) = 2.503, p = 0.008, Cohen’s d = 0.688, see Figure 2).

**Figure 2.**
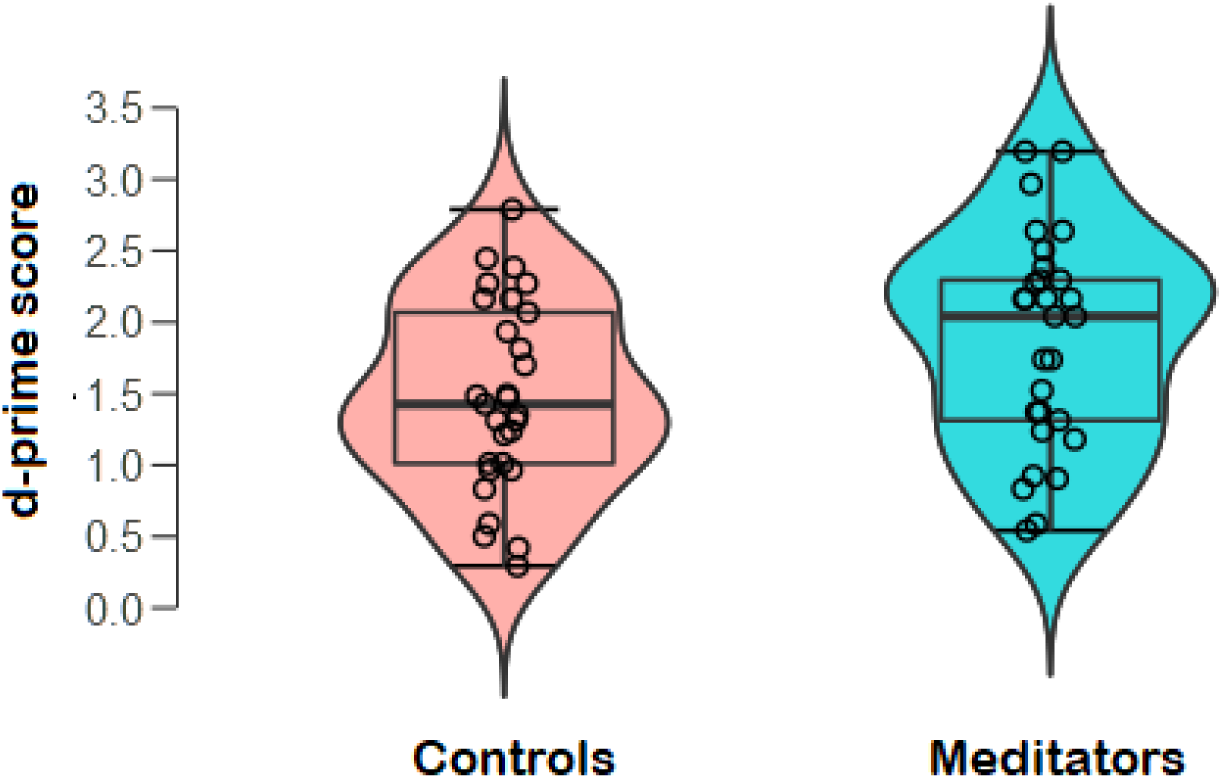
Box and violin plot of d-prime scores from the modified Sternberg task for each group (p = 0.008, Cohen’s d = 0.688). Circles reflect scores from each individual and the outer curve reflects the distribution of scores in each group.

There were no group differences in reaction time (F(1,56) = 0.118, p = 0.732, η^2^= 0.002), nor a group by probe condition interaction (F(1,56) = 1.464, p = 0.231, η^2^= 0.025), nor was there a difference in probe condition reaction time (F(1,56) = 1.188, p = 0.280, η = 0.020, η_p_^2^ = 0.021). The lack of group differences in reaction time suggests that the meditation group did not obtain higher accuracy scores simply due to a speed / accuracy trade off. Means and standard deviations are shown in table 2. A significant main effect of group in the number of accepted epochs was detected with the meditation group showing a higher number of epochs (F(1,56) = 8.506, p = 0.005, η^2^ = 0.132). Because differences in the number of epochs included in ERP averages can affect comparisons, post-hoc validation comparisons were conducted between the groups after a random selection of epochs were removed from the meditation participants until each meditation participant had an equal number of epochs to a control participant. This analysis showed the same results as the main comparison, and is reported in the supplementary materials.

**Table 2.**
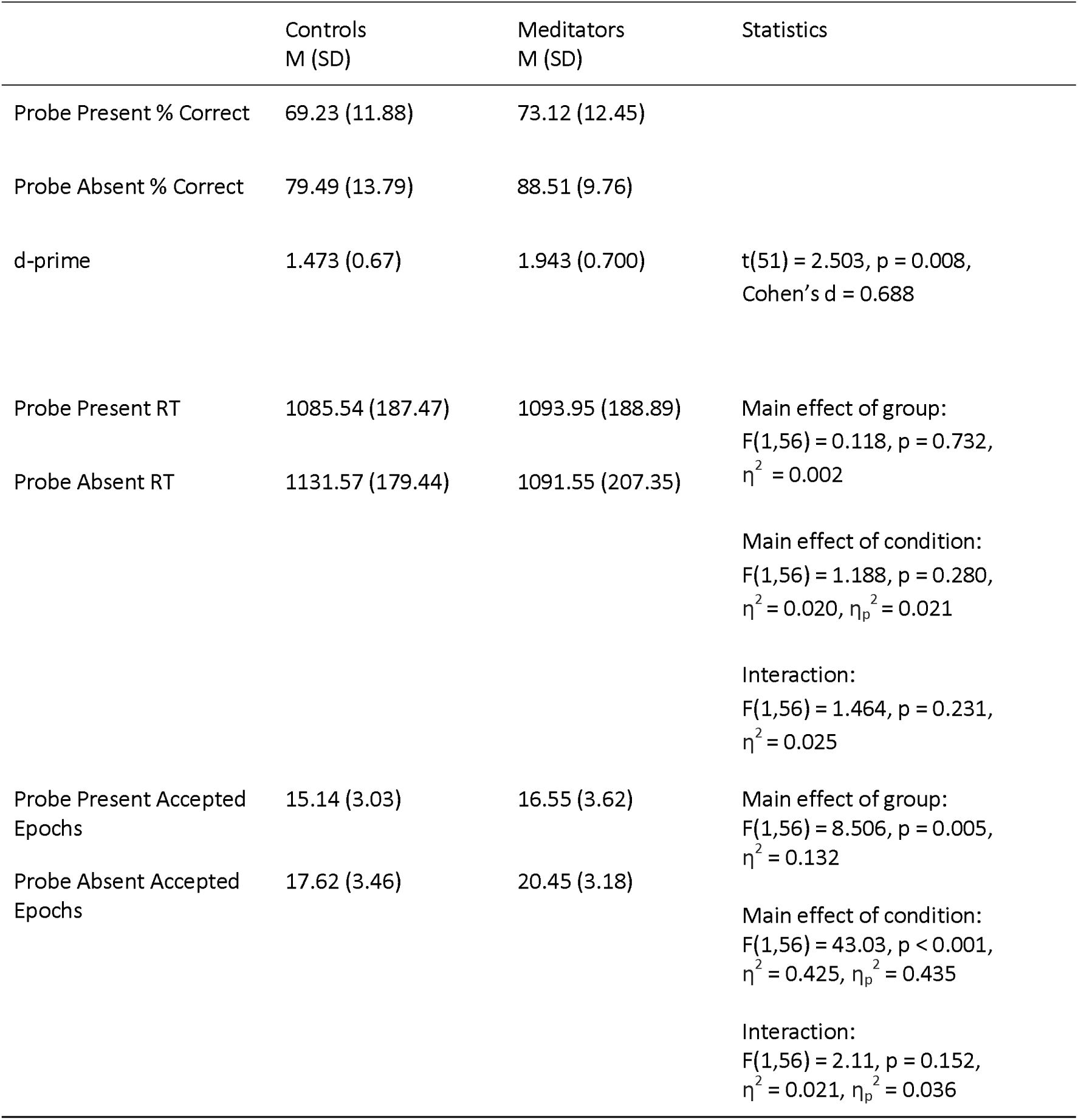
Accuracy, reaction time and accepted epoch data for the two groups

### WM delay period BLC alpha, theta, and 1/f aperiodic activity comparisons

No significant group differences were present for either alpha BLC power in the WM delay period in RMS or TANOVA comparisons across the entire epoch using RAGU (all FDR-p and p > 0.05). Similarly, no significant group differences were present for theta BLC power in the WM delay period in RMS or TANOVA comparisons across the entire epoch using RAGU (all FDR-p and p > 0.05). When 1/f aperiodic activity was separated from the oscillatory activity, the meditation group showed a higher proportion of participants with theta peaks at Fz (Chi squared = 5.156, p = 0.0232) and a trend towards an interaction between group and electrode for 1/f aperiodic offset at parieto-occipital electrodes (F(1,56) = 4.025, p = 0.050, η^2^ = 0.057, η_p_^2^ = 0.067), with meditators showing a larger difference between the two electrodes than controls (with lower 1/f aperiodic offset values at PO7). The 1/f aperiodic slope showed no differences between the groups or interactions between group and electrode, and there was no significant main effect of 1/f aperiodic offset at parietal electrodes or at Fz (all p > 0.05, statistics reported in table 3). No significant differences were detected in alpha centre frequency, bandwidth, or amplitude across PO7 and PO8 (all p > 0.05). Nor were any significant differences detected in theta centre frequency, bandwidth, or amplitude at Fz (all p > 0.05). Values and statistics are reported in table 3.

**Table 3.**
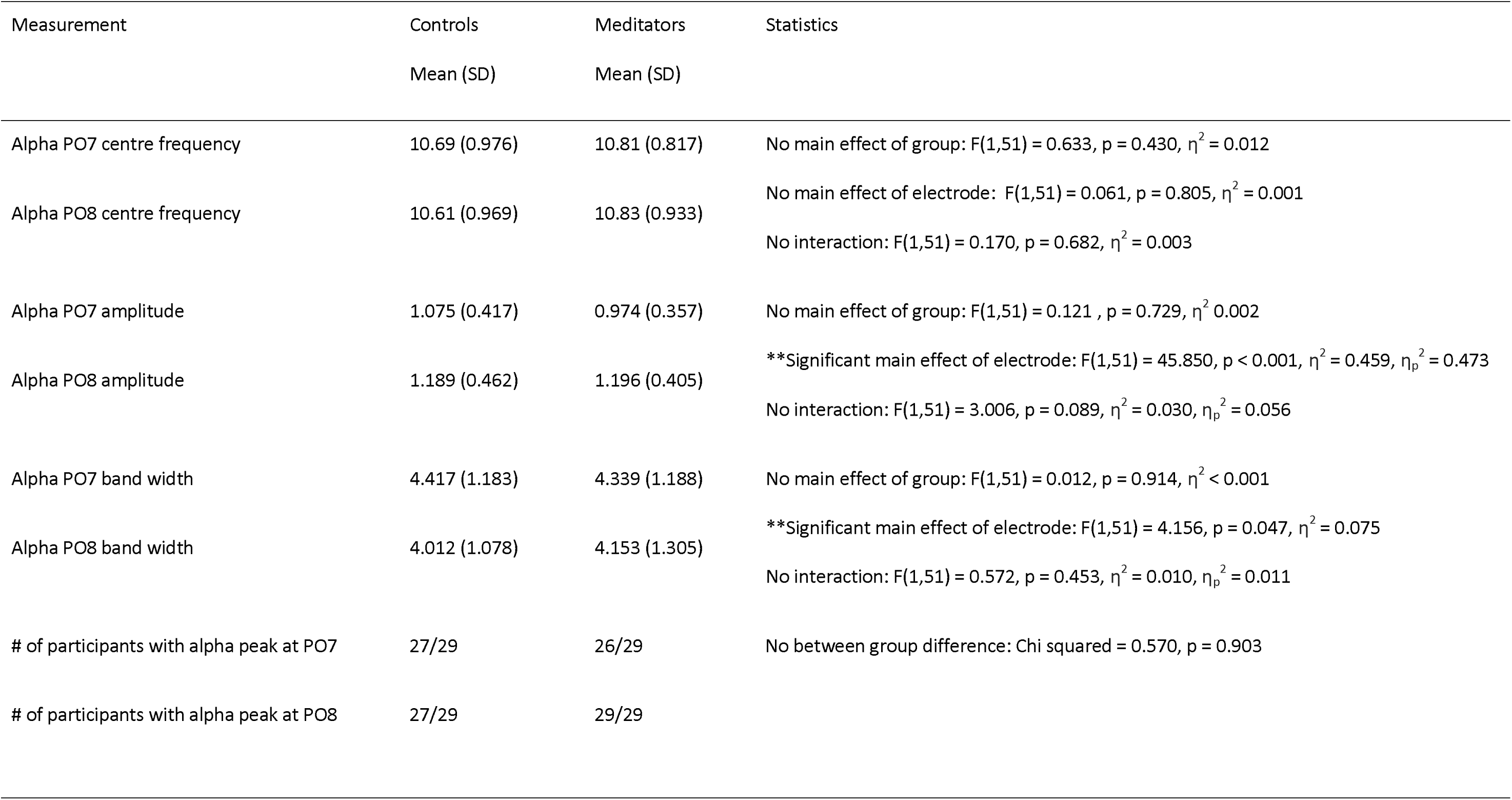
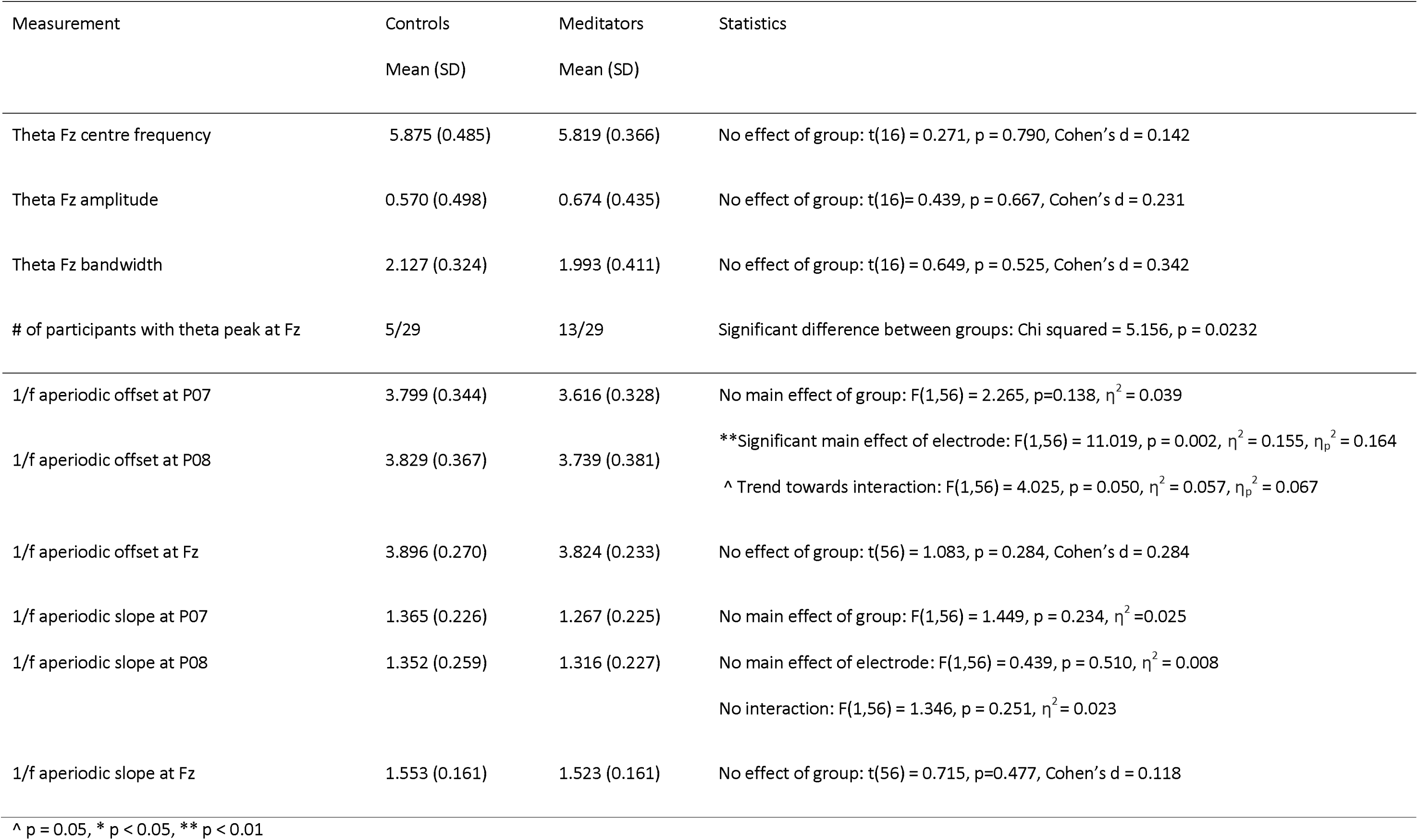
WM delay period 1/f aperiodic activity statistics and oscillation statistics when measured independently of the 1/f aperiodic activity

| Measurement | Controls | Meditators | Statistics |
| --- | --- | --- | --- |
|  | Mean (SD) | Mean (SD) |  |
| Alpha PO7 centre frequency | 10.69 (0.976) | 10.81 (0.817) | No main effect of group: $F(1,51) = 0.633$, $p = 0.430$, $\eta^2 = 0.012$ |
| Alpha PO8 centre frequency | 10.61 (0.969) | 10.83 (0.933) | No main effect of electrode: $F(1,51) = 0.061$, $p = 0.805$, $\eta^2 = 0.001$<br>No interaction: $F(1,51) = 0.170$, $p = 0.682$, $\eta^2 = 0.003$ |
| Alpha PO7 amplitude | 1.075 (0.417) | 0.974 (0.357) | No main effect of group: $F(1,51) = 0.121$, $p = 0.729$, $\eta^2 = 0.002$ |
| Alpha PO8 amplitude | 1.189 (0.462) | 1.196 (0.405) | **Significant main effect of electrode: $F(1,51) = 45.850$, $p < 0.001$, $\eta^2 = 0.459$, $\eta_p^2 = 0.473$<br>No interaction: $F(1,51) = 3.006$, $p = 0.089$, $\eta^2 = 0.030$, $\eta_p^2 = 0.056$ |
| Alpha PO7 band width | 4.417 (1.183) | 4.339 (1.188) | No main effect of group: $F(1,51) = 0.012$, $p = 0.914$, $\eta^2 < 0.001$ |
| Alpha PO8 band width | 4.012 (1.078) | 4.153 (1.305) | **Significant main effect of electrode: $F(1,51) = 4.156$, $p = 0.047$, $\eta^2 = 0.075$<br>No interaction: $F(1,51) = 0.572$, $p = 0.453$, $\eta^2 = 0.010$, $\eta_p^2 = 0.011$ |
| # of participants with alpha peak at PO7 | 27/29 | 26/29 | No between group difference: Chi squared = 0.570, $p = 0.903$ |
| # of participants with alpha peak at PO8 | 27/29 | 29/29 |  |
|  | Mean (SD) | Mean (SD) |  |
| Theta Fz centre frequency | 5.875 (0.485) | 5.819 (0.366) | No effect of group: $t(16) = 0.271$, $p = 0.790$, Cohen's $d = 0.142$ |
| Theta Fz amplitude | 0.570 (0.498) | 0.674 (0.435) | No effect of group: $t(16) = 0.439$, $p = 0.667$, Cohen's $d = 0.231$ |
| Theta Fz bandwidth | 2.127 (0.324) | 1.993 (0.411) | No effect of group: $t(16) = 0.649$, $p = 0.525$, Cohen's $d = 0.342$ |
| # of participants with theta peak at Fz | 5/29 | 13/29 | Significant difference between groups: Chi squared = 5.156, $p = 0.0232$ |
| 1/f aperiodic offset at P07 | 3.799 (0.344) | 3.616 (0.328) | No main effect of group: $F(1,56) = 2.265$, $p = 0.138$, $\eta^2 = 0.039$ |
| 1/f aperiodic offset at P08 | 3.829 (0.367) | 3.739 (0.381) | **Significant main effect of electrode: $F(1,56) = 11.019$, $p = 0.002$, $\eta^2 = 0.155$, $\eta_p^2 = 0.164$<br>^ Trend towards interaction: $F(1,56) = 4.025$, $p = 0.050$, $\eta^2 = 0.057$, $\eta_p^2 = 0.067$ |
| 1/f aperiodic offset at Fz | 3.896 (0.270) | 3.824 (0.233) | No effect of group: $t(56) = 1.083$, $p = 0.284$, Cohen's $d = 0.284$ |
| 1/f aperiodic slope at P07 | 1.365 (0.226) | 1.267 (0.225) | No main effect of group: $F(1,56) = 1.449$, $p = 0.234$, $\eta^2 = 0.025$ |
| 1/f aperiodic slope at P08 | 1.352 (0.259) | 1.316 (0.227) | No main effect of electrode: $F(1,56) = 0.439$, $p = 0.510$, $\eta^2 = 0.008$<br>No interaction: $F(1,56) = 1.346$, $p = 0.251$, $\eta^2 = 0.023$ |
| 1/f aperiodic slope at Fz | 1.553 (0.161) | 1.523 (0.161) | No effect of group: $t(56) = 0.715$, $p = 0.477$, Cohen's $d = 0.118$ |

^ $p = 0.05$ , \* $p < 0.05$ , \*\* $p < 0.01$

### ERP Comparisons During WM Probe Period

The TCT showed significant signal indicating consistency of neural activity within all groups across the entire epoch except during the period prior to the stimulus to 80 ms after probe stimulus presentation (see supplementary materials Figure S1).

### WM probe period GFP test

To compare the strength of neural responses in the WM probe period between groups we conducted a GFP test. A significant difference that survived duration multiple controls (but not global count multiple controls) was present between groups from 399 to 481 ms, and 489 to 585 ms, with meditators showing reduced GFP values in both these time windows (duration control = 55 ms, global count p = 0.0512, global count FDR-p = 0.1536). When averaged across both significant periods, including the break in the middle, the effect was significant (averaged p = 0.0062, averaged FDR-p = 0.0098, η^2^ = 0.1251, see Figure 3). Across both groups, averaged GFP amplitude during this time period did not covary with d-prime scores (p = 0.9222). Nor did GFP amplitude covary with d-prime scores within just the meditation group (p = 0.1296), nor the control group (p = 0.9110). Because the groups showed differences in the number of accepted epochs included in the analyses, a validation analysis was performed with number of included epochs matched between the groups. Additionally, although probe present and probe absent conditions were averaged to ensure enough epochs for valid ERP comparisons in the main analysis, exploratory analyses were conducted separating the two conditions. These two analyses are reported in the supplementary materials. Both analyses showed similar results to the main analysis, with a significant main effect of group showing the meditation group with smaller GFP values (but no interaction between group and probe present / probe absent). However, the significant time period was shorter in the matched epoch analysis (however, averaged across the significant time window from the main analysis, the matched epoch was still significant, p = 0.0218). Both the time window of the GFP difference and the topography of activity in this time window (see the microstate analysis, Figure S4) indicated that the groups differed in overall amplitude of the parietal old/new effect component.

**Figure 3.**
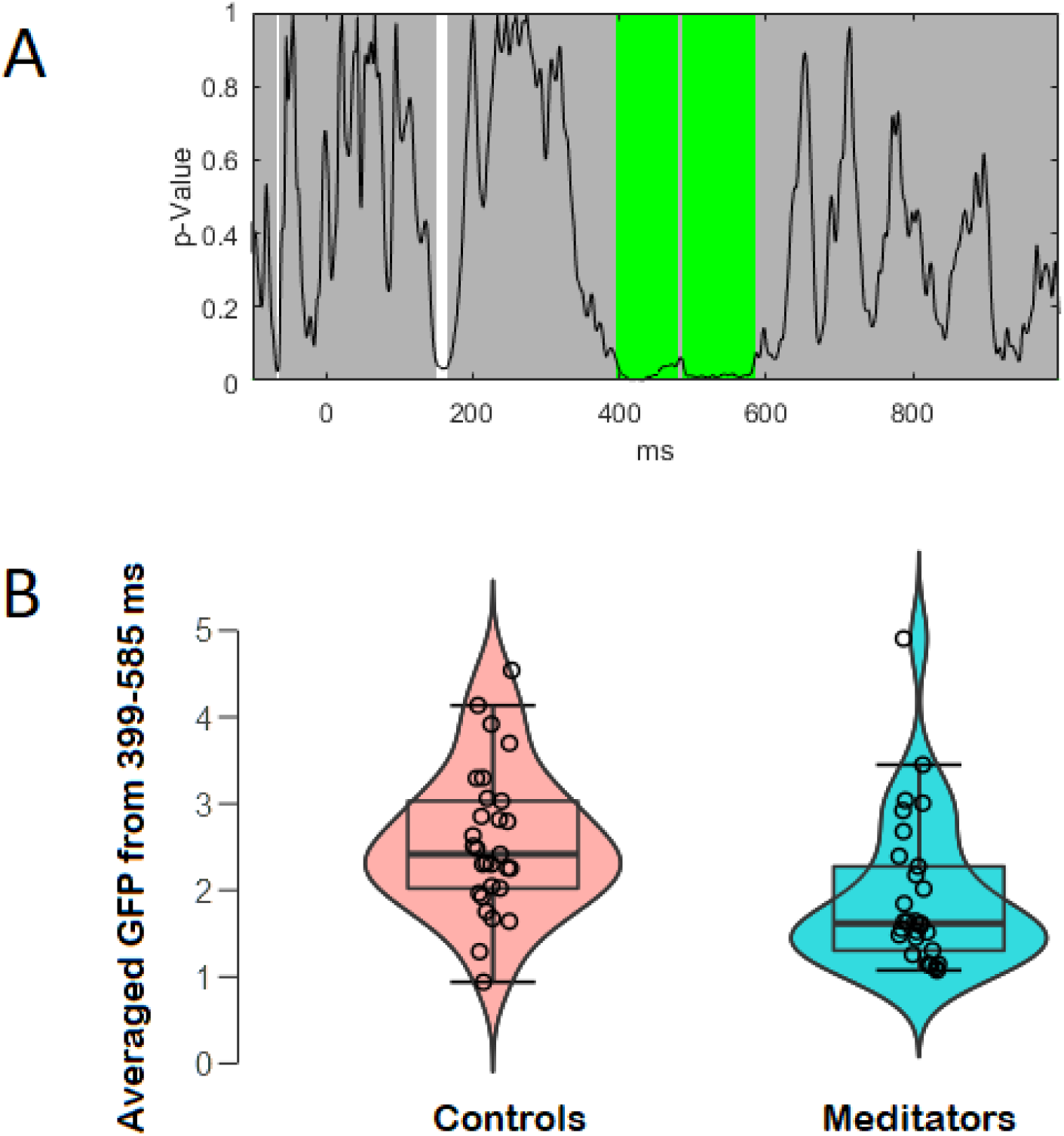
Significant between group difference in GFP from 399 to 585 ms following the WM probe onset. A – p-values of the between group comparison for the real data against 5000 randomly shuffled permutations across the entire epoch (green periods reflect periods that exceed the duration control for multiple comparisons across time = 55 ms, averaged across the time period of significance p = 0.0062, FDR-p = 0.0098, η^2^= 0.1251, global count p = 0.0512, global count FDR-p = 0.1536). B – Box and violin plot of the averaged GFP across the entire window of significance (from 399 to 585 ms). Circles reflect scores from each individual and the outer curve reflects the distribution of scores in each group.

### WM Probe Period TANOVA

The TANOVA showed a significant difference between the two groups in the distribution of neural activity, which lasted from 294 to 672 ms following the probe stimuli (see Figure 4). The significant difference survived all multiple comparison controls (duration control = 49 ms, global count p = 0.0028, global count FDR-p = 0.0168, averaged across significant time period p = 0.002, FDR-p = 0.0186, η^2^= 0.0903). The difference appears to be driven by more right frontal positivity in meditators at the beginning of the significant period, which shifted towards more fronto-central positivity towards the end of the significant period. Similar to the GFP comparisons, validation checks were conducted to control for differences in the number of accepted epochs included in analyses (reported in the supplementary materials). Both analyses with matched numbers of epochs across the groups and analyses of probe present / absent conditions separated showed similar results to the main analysis, with a significant main effect of group during the same time period with the same pattern. No interaction was present between group and probe present / probe absent conditions (all p > 0.05). The 294 to 672 ms period of significance was separated into three separate time periods based on distinct microstates for analysis of the relationship between d-prime and neural activity (310-425 ms, 425-600 ms and 600-672 ms). None of these time periods contained neural activity that showed a relationship to d-prime scores (all p > 0.05, reported in more detail in the supplementary materials).

**Figure 4.**
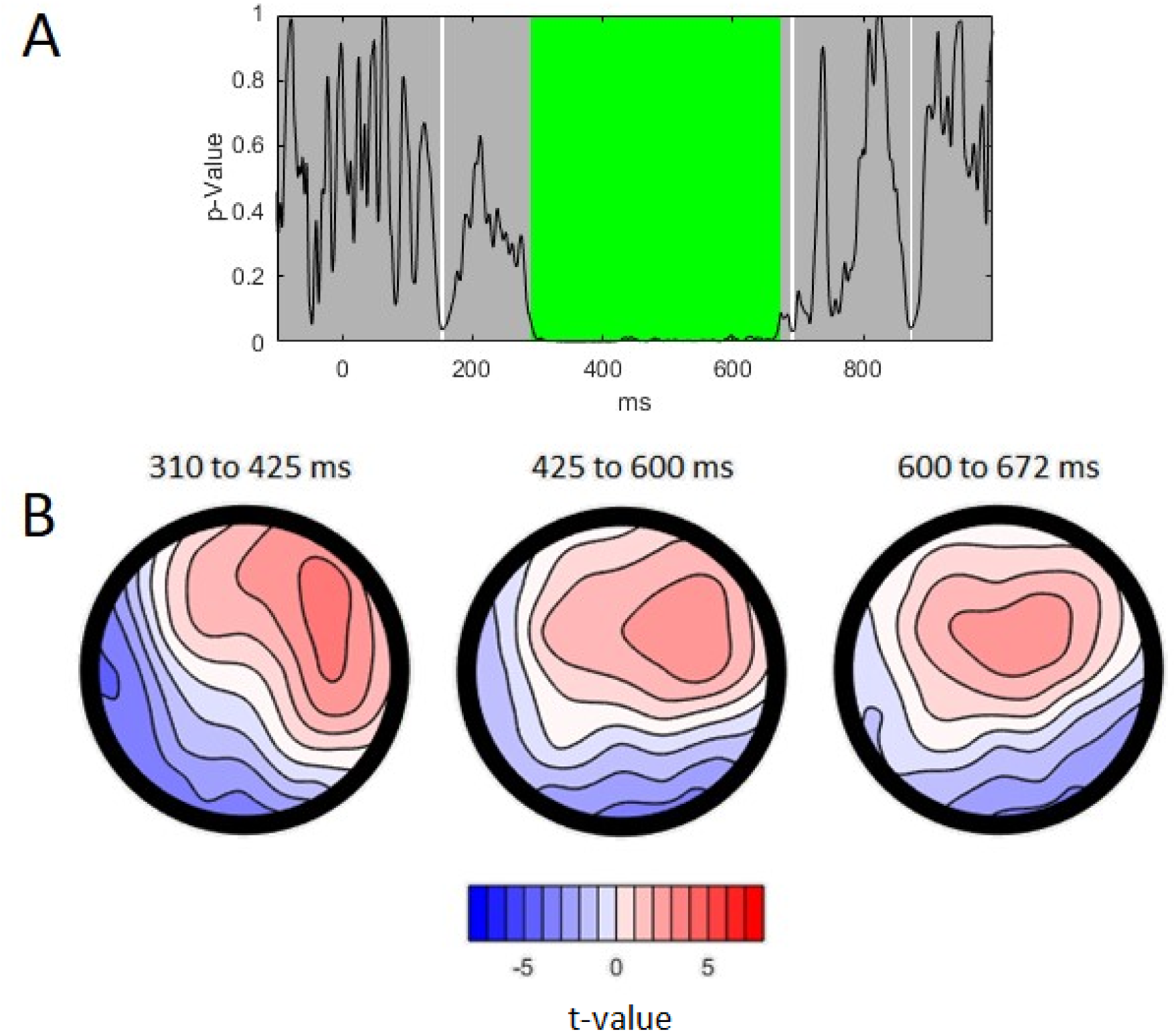
TANOVA main group effect from 294 to 672 ms. A – p-values of the between-group comparison for the real data against 5000 randomly shuffled permutations across the entire epoch (global count p = 0.0028, global count FDR-p = 0.0168, green periods reflect periods that exceed the duration control for multiple comparisons across time, global duration control = 49 ms, averaged across the significant window p = 0.002, η^2^ = 0.0903). B - t-maps for meditators topography minus control topography during the significant time window, split into periods reflecting distinct microstates as reported below (for the average activity over the first time period, η^2^ = 0.1104, for the average activity over the second time period η = 0.0781, and for the average activity over the third time period η^2^ = 0.0692).

### Exploratory source analysis of the WM probe period

Topographies that differed between the groups in the TANOVA comparisons reflect differences in recruitment of underlying generators of neural activity (Koenig et al., 2011), so source analyses were used to characterise which generators were likely to differ between groups. The microstate analysis identified three separate time windows that depicted distinct topographies during the time window that groups showed significant differences in TANOVA comparisons. These time windows were from 310 to 425 ms (reflecting the FN400, see Figure 5), 425 to 600 ms, and 600 to 672 ms (both reflecting the parietal old/new effect, see Figures 6 and 7). The supplementary materials report how decisions for these time periods were made.

**Figure 5.**
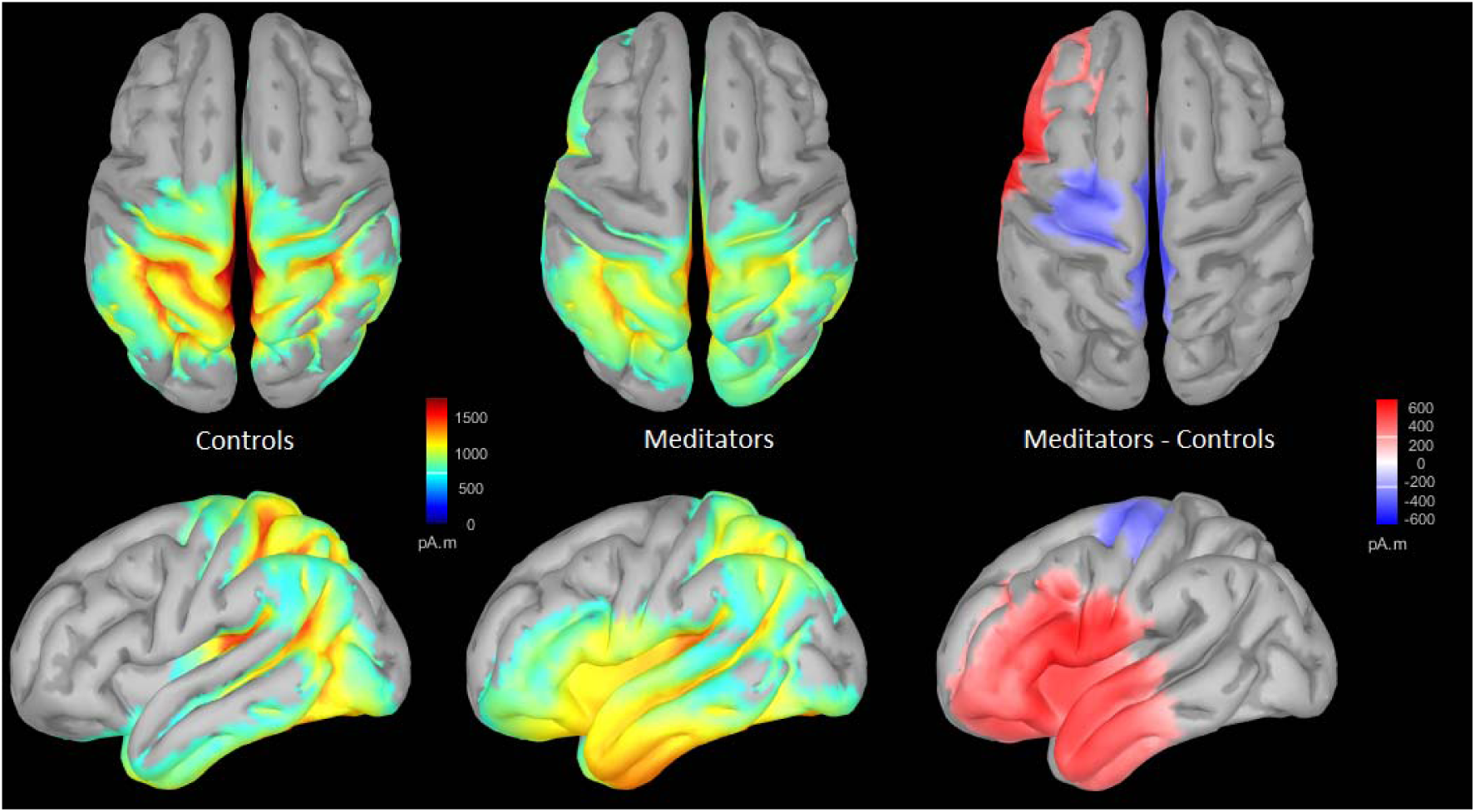
Source reconstruction during the 310 to 425 ms window using sLORETA and minimum norm imaging, unconstrained to cortex (to minimise assumptions). Activity during this window is likely to reflect the FN400. Note that the source modelling method used depicts only absolute activation so that more positive values in source analysis reflect stronger values in either positive or negative directions. Difference maps reflect meditator minus control activity (red reflecting more activity in meditators compared to controls, blue reflecting less activity in meditators).

**Figure 6.**
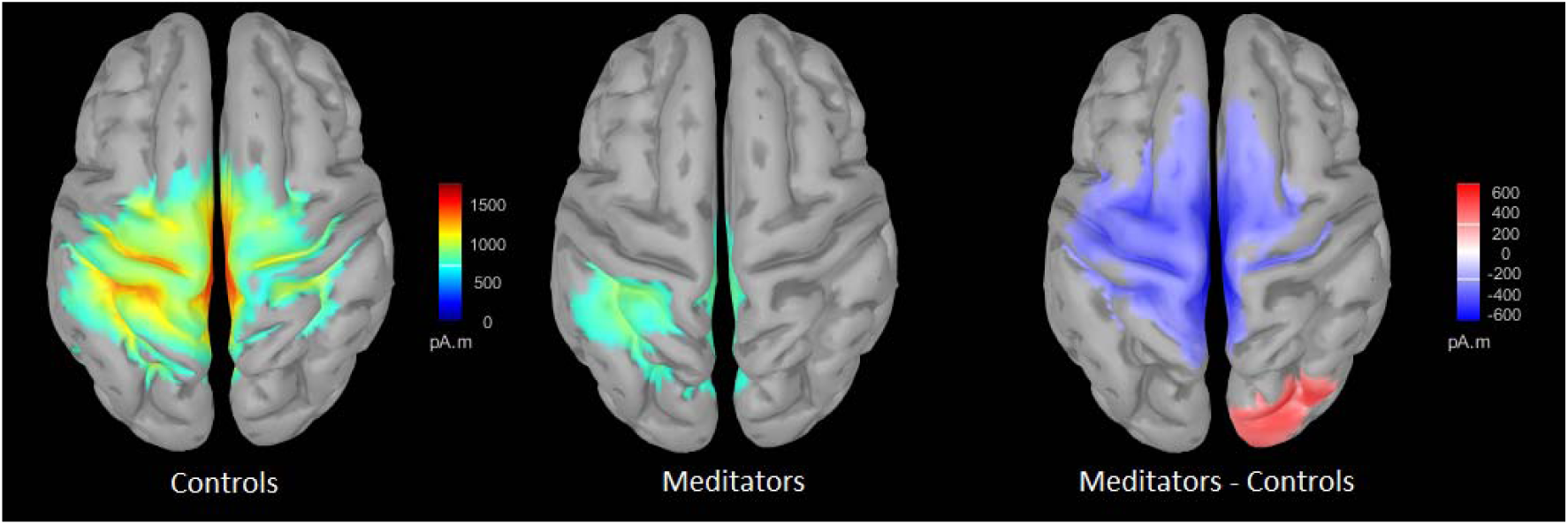
Source reconstruction during the 425 to 600 ms window using sLORETA and minimum norm imaging, unconstrained to cortex (to minimise assumptions). Activity during this window is likely to reflect the parietal old/new effect. Note that the source modelling method used depicts only absolute activation so that more positive values in source analysis reflect stronger values in either positive or negative directions. Difference maps reflect meditator minus control activity (red reflecting more activity in meditators compared to controls, blue reflecting less activity in meditators).

**Figure 7.**
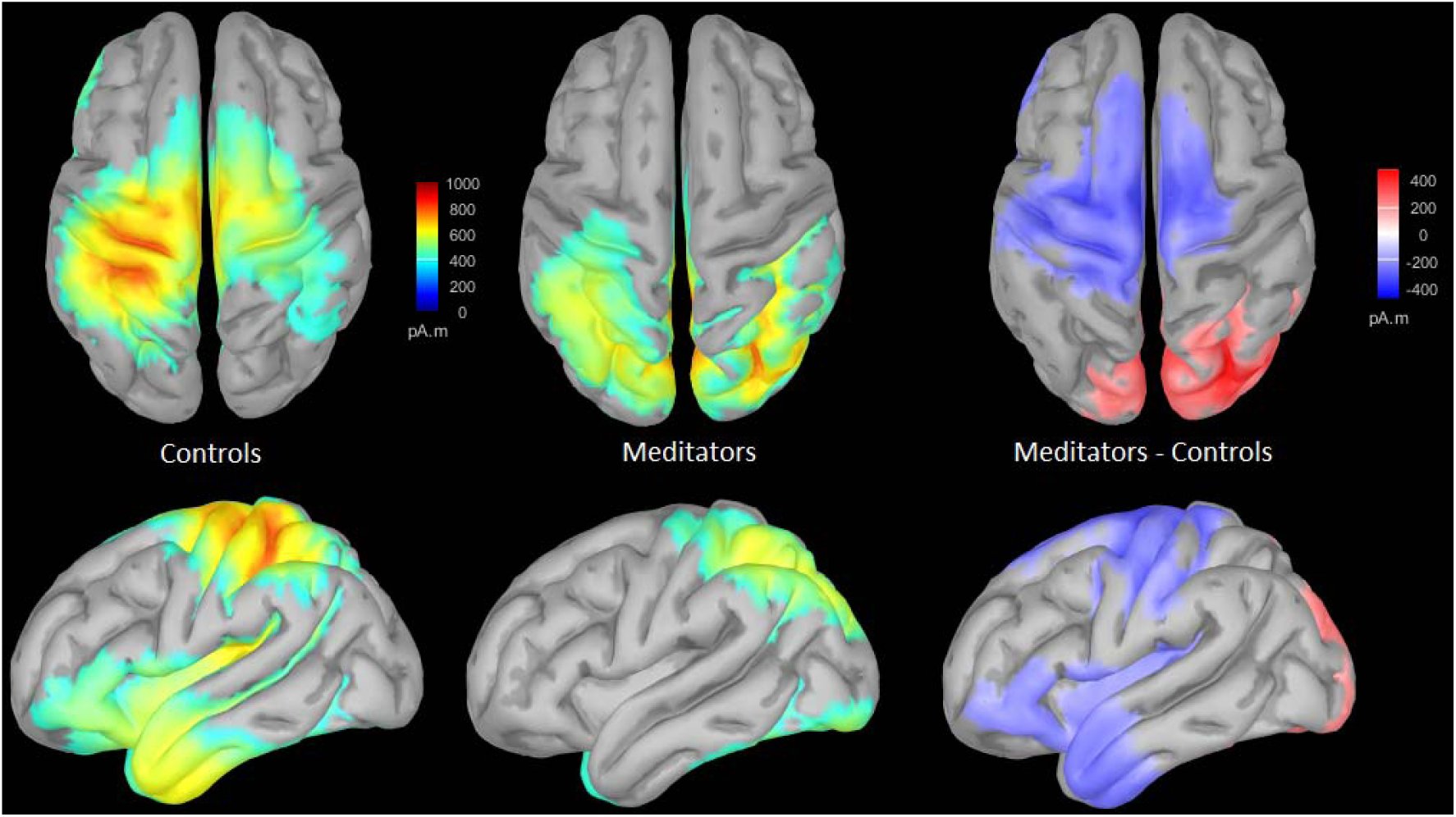
Source reconstruction during the 600 to 672 ms window using sLORETA and minimum norm imaging, unconstrained to cortex (to minimise assumptions). Activity during this window is likely to reflect the parietal old/new effect. Note that the source modelling method used depicts only absolute activation so that more positive values in source analysis reflect stronger values in either positive or negative directions. Difference maps reflect meditator minus control activity (red reflecting more activity in meditators compared to controls, blue reflecting less activity in meditators).

Source analysis indicated that from 310 to 425 ms post stimulus, both groups showed broad parietal activation, and the meditation group additionally showed left temporal activation (see Figure 6). Absolute activations are depicted on the cortex from source analyses in Figures 6,7, and 8. Note that the source modelling method used depicts absolute activation, so that more positive values in source analysis reflect stronger values in either positive or negative voltages. As such, the increased left temporal activation in meditators in the 310 to 425 ms period is likely to reflect more negative activity in the left temporal regions at the scalp level (as can be seen in the TANOVA comparisons). From 425 to 600 ms, both groups showed centro-parietal activation, although the meditation group showed a much smaller region of activation that exceeded the threshold (see Figure 7). This is likely to reflect the reduced overall neural response strength shown in the GFP test, as the source analysis does not separate differences in the distribution of activity from differences in amplitude (in contrast to analyses performed with RAGU). Lastly in the 600 to 672 ms window, meditators showed less left temporal and fronto-central activation, and more occipital activation than the control group, probably reflecting a combination of differences in overall neural response strength and in the distribution of activity (see Figure 8). Taken together, source analysis of these three time windows suggests that meditators activate their left temporal region earlier than controls, and that this activity is completed earlier, with less left temporal activity and less overall activity later in the epoch.

## Discussion

This study cross-sectionally compared neural activity related to working memory in long term mindfulness meditators and demographically matched non-meditating controls. The meditation group showed higher accuracy in the WM task. Concurrent with this higher accuracy, the meditation group showed an altered distribution of both the FN400 and parietal old/new effect neural activity during the WM probe period, as well as a probable reduction of overall neural response strength during parietal old/new effect (with medium effect sizes). Scalp analysis of the EEG data suggested that the meditation group showed more left temporal negative voltages and frontal right positive voltages during the FN400. During the parietal old/new effect the meditation group showed more fronto-central positive voltages and less positive parietal voltages. Source analysis suggested the differences were the result of more neural activity in the left temporal lobe in the meditation group during the FN400, followed by less neural activity in central-parietal regions during the parietal old/new effect. The meditation group showed a higher proportion of participants with fronto-midline theta peaks above the 1/f aperiodic activity during the WM delay period. However, no overall differences were detected in WM delay period neural activity, including measures of theta, alpha, and 1/f aperiodic activity (although there was a trend towards interaction between group and electrode for 1/f aperiodic offset at parieto-occipital electrodes, with the meditation group showing lower values at PO7 only).

### Meditators showed altered distribution and probable reduced amplitude of WM probe period ERP

The most conservative interpretation of the WM probe period ERP differences is that meditators show altered distributions of neural activity and reduced neural response strengths during the recall period. Further interpretation involves engaging in reverse inference – ascribing differences between groups as reflecting differences in functional processes. Conclusions drawn from reverse inference are not deductively valid (particularly when activated regions are broad as per EEG source analyses) as each brain region (and each ERP component) has multiple functional associations (Poldrack, 2006). As such, increases or decreases in specific brain regions (or ERP components) could reflect differences in a range of possible functions. Interpretation is also complicated by the complexity of the current results (reflecting differences in multiple WM ERP components as well as both distribution and overall neural response strength differences). However, with these caveats in mind, reverse inference can still provide information (Poldrack, 2006), particularly when the task generating the activity is taken into account (Hutzler, 2014).

With regards to the differences in the current study, WM probe period ERPs in the modified Sternberg task have been suggested to reflect retrieval, and are related to memory scanning and decision making (Ergen et al., 2012). WM probe period ERPs similar to those detected in our study have also been suggested to reflect a process that functions to inhibit widespread cortical regions, suppressing irrelevant neural activity from interrupting the WM relevant information (W Klimesch et al., 2000). In memory tasks more generally, the FN400 has been suggested to reflect familiarity related processing (Curran & Cleary, 2003; Duarte et al., 2004), conceptual priming (Woodruff et al., 2006), and semantic processing (binding stimulus information to recalled information) (Federmeier & Laszlo, 2009; Yonelinas, 2002). FN400 activity is thought by some research to be generated in part by the left temporal region, and activation of this region during the FN400 is particularly related to attention mediated familiarity processing, with more negative activity in response to stimuli that are not familiar (Stróżak et al., 2016). Activity in the left temporal region is also associated with phonological processing in WM tasks (Vigneau et al., 2006), and left hemisphere effects are also more prominent for verbal WM tasks (Nagel, Herting, Maxwell, Bruno, & Fair, 2013; Osaka, Komori, Morishita, & Osaka, 2007). Based on this interpretation, the increased negativity in the temporal region in meditators could suggest they were less familiar with the probe stimuli. However, a reduction in familiarity related processing in probe present trials would likely lead to reduced performance in those trials. Neither an interaction between group and trial type, nor reduced performance in probe present trials in meditators was found. As such, it seems more likely that the increased temporal negativity in meditators reflects increased conceptual, semantic or phonological processing. As well as the left temporal activity in the FN400, positive voltages in right frontal regions have been suggested to reflect general monitoring and decisional processes thought to be generated by the right DLPFC (Hayama, Johnson, & Rugg, 2008). As such, the more right frontal positive voltage distribution in the meditation group might suggest they engage in more general monitoring and decision processes. More frontal distributions of the P3 have also been found in our previous research in meditators (N. W. Bailey et al., 2019; Wang et al., 2019), and we have previously suggested these alterations reflect enhanced attention.

As well as differences in the FN400, our results showed differences in the distribution and probable differences in the amplitude of the parietal old/new effect. The parietal old/new ERP following the FN400 has been associated with conscious recollection, with its amplitude reflecting attention orientation to recalled information (Duarte et al., 2004; Rugg & Curran, 2007; Rugg & Henson, 2002; Wagner et al., 2005; Woodruff et al., 2006; Yonelinas, 2002). In particular, the left parieto-occipital positivity (activity of which the meditation group showed lower amplitudes) is larger when items are consciously remembered and during successful memory performance (Friedman & Johnson Jr, 2000). Similar to the alterations in the FN400, the reduced left parieto-occipital positive voltage and reduced overall amplitude in the meditation group could then be expected to have resulted in lower performance, which was not the case. It may be that the meditation group performed the functions associated with the parietal old/new effect in other regions or more efficiently, leading to a reduction in the ERP without a reduction in performance.

While the distribution differences passed all multiple comparison controls (and as such likely represent true differences between the groups), the reduction in neural response strength was significant when using multiple comparison control for the duration of the effect, but not when controlling for all timepoints measured in the epoch. Future analyses focusing specifically on the significant window from our research are likely to show differences between groups, but our exploratory approach including the entire epoch cannot draw firm conclusions. The medium-to-large effect size of the GFP difference suggests the finding is likely to reflect a true difference present in the two populations sampled, particularly since global control multiple comparisons used in the current study is considered overly conservative (Koenig et al., 2011), and controls for the number of significant time points in the real data but do not consider the strength of significant differences.

Together, these differences suggest that rather than a simple increase in strength of the effective WM strategies that high performing non-meditator participants have shown in previous research (Maurer et al., 2015; Scheeringa et al., 2009), the meditation group used a different strategy, engaging different neural processes and thus showing an altered distribution of neural activity (and potentially a reduced overall amplitude). These differences might reflect altered neural processes in the meditation group, whereby frontally driven decision relevant processes might be activated earlier, allowing meditators more processing time before responding, and thus enabling more accuracy with less overall neural response strength. In support of this explanation, the meditation group’s GFP amplitude peaked at about 300 ms after the probe while the control group’s GFP amplitude continued to increase until a larger and later peak at 400 ms (see microstate analyses in supplementary materials). While not specifically measured in the current study, it is worth noting that the P3 latency is thought to reflect processing speed (Pontifex et al., 2009). It should be emphasised, however, that none of these differences in neural activity were related to accuracy (a finding that is explored in a later section), and that while this explanation does fit the data, it is simply a possibility until further research explores the functional relevance of these differences in neural activity in more detail. As such, these interpretations should be considered as tentative, and further research is required to eliminate other potential explanations as well as replicate the current results before these explanations could be considered accurate.

### Lack of relationships between neural activity differences between groups and performance measures

Across the different ERP components that showed differences in neural activity, only the early component of the FN400 showed what could be reported as a trend towards a relationship between d-prime scores and the distribution of neural activity (p = 0.0672). This result was obtained assessing the relationship across all participants, and as such may be confounded by between group differences in both neural activity and behavioural performance. Thus, it seems unlikely that any particular measure of altered neural activity in meditators that we detected is the explanation for their improved WM performance. A number of potential interpretations for this are possible. The first (and in our view most likely) is that the improved WM performance is related to differences in neural processes that are causal for the differences in the measures we have detected, so that if we were to measure these deeper mechanisms (that lead to both the differences in the recall ERP and the improved performance) we would find a relationship to WM performance. The second potential explanation is that the differences in neural activity we detected do not relate to WM performance per se, but instead relate to differences in the experience of performing a WM task between the groups (perhaps differences in attention or non-judgemental acceptance that do not convert to differences in WM performance). If this explanation is true, we suggest that another (not yet detected) difference in neural activity between the groups is required to explain the differences in WM performance. Finally, it may be that the relationship between WM performance and these neural measures is non-linear, or that the relationships are weak and require larger sample sizes to detect.

### No differences in WM delay period theta, alpha, or 1/f aperiodic activity

Although research has suggested WM delay period parieto-occipital alpha and fronto-midline theta to be related to WM performance (Maurer et al., 2015; Scheeringa et al., 2009), differences in the BLC power, amplitude, centre frequency, or bandwidth of these oscillations were not present in the meditation group (both when measured by traditional power computations across all electrodes, and measured independently of the 1/f aperiodic activity at specific electrodes of interest). As such, differences in these oscillations do not seem to be the explanation for the enhanced performance in the meditation group. However, meditators did show a higher proportion of participants with fronto-midline theta peaks above the 1/f aperiodic activity. Theta activity has been related to attentional function, particularly cognitive control (Cavanagh & Frank, 2014; Rac-Lubashevsky & Kessler, 2018; Sauseng et al., 2010; Sauseng et al., 2007), and previous research has indicated that meditators show increases in theta activity (Kerr et al., 2013; Lagopoulos et al., 2009; Tang et al., 2009). As such, it may be that meditation practice enabled a higher proportion of the meditation group to generate theta oscillations to maintain attention on the task during the WM delay period. Having said that, it is also worth noting that when separated from the aperiodic signal, only a minority of participants in either group showed theta peaks during the WM delay period (18/58 in total). Previous research has suggested that theta activity is functionally relevant in the Sternberg task, but until now this has not been tested with the 1/f aperiodic activity removed from oscillation measurements (Brookes et al., 2011; O. Jensen & Tesche, 2002; Kottlow et al., 2015; Maurer et al., 2015; L. Payne & Kounios, 2009; Scheeringa et al., 2009). This finding emphasises that it will be important for future research exploring the functional importance of different frequencies to measure oscillatory activity separately from 1/f aperiodic activity.

Similarly, no differences were detected in 1/f aperiodic slope. 1/f aperiodic activity is generated by population spiking synchrony, which has been suggested to reflect excitation / inhibition balances (Peterson et al., 2018). These decreases or increases in proportions of inhibition and excitation could reflect alterations in numbers of the different types of neurons, or alterations in the strength of synaptic connections (Gao et al., 2017). Although the current results suggest that 1/f aperiodic activity measures do not differ in meditators, this is the first exploration of 1/f aperiodic activity in meditators, and the analyses were only post-hoc exploratory analyses restricted to three electrodes in a single task. Further research is required before the current null result could be confirmed. It would also be interesting to compare meditators and controls in 1/f aperiodic activity in other tasks or during the meditation state. This research is worth pursuing, as the altered inhibition / excitation balances have been suggested as potential explanations underpinning the benefits of meditation (Edwards, Peres, Monti, & Newberg, 2012; Guglietti, Daskalakis, Radhu, Fitzgerald, & Ritvo, 2013).

In contrast to the null results for alpha, theta, and 1/f aperiodic slope, there was a trend towards an interaction between group and electrode for 1/f aperiodic offset at parieto-occipital electrodes, with meditators showing lower values at PO7 only. Offset is positively related to overall neuronal firing rates / spiking activity (Manning et al., 2009; Miller et al., 2012), and the fMRI bold signal (Winawer et al., 2013). This might reflect meditators showing less overall neural activity in the left parieto-occipital region during the retention period. However, given the exploratory nature of the 1/f aperiodic analyses, the number of uncontrolled multiple comparisons, and the trend level finding, we are hesitant to draw conclusions from this finding, and are uncertain if the result would replicate. Further research exploring 1/f aperiodic metrics in samples of experienced meditators is required.

### Strengths, Limitations and Future Directions

Although the control group in our study was well-matched to the meditation group in terms of age, gender and years of education, the primary limitation of cross-sectional research is that it does not offer any information about causation. Longitudinal studies with active control comparison groups have suggested that mindfulness meditation is causal in WM performance improvements (Jha et al., 2010; Mrazek et al., 2013; Quach et al., 2016; Van Vugt & Jha, 2011; Zeidan et al., 2010). Similarly, reduced neural activity related to distractor flickering in stimuli in WM tasks has been shown to result from mindfulness training (Schöne, Gruber, Graetz, Bernhof, & Malinowski, 2018). While these results suggest that meditation may be causal in the current results, we cannot rule out the possibility that self-selection biases in those who choose to meditate underpin the differences in neural activity related to WM in the current study. As such, the most parsimonious and robust interpretation of the results of cross-sectional studies of experienced meditators is that the differences relate to “leading a life that involves meditation” (N. W. Bailey et al., 2019).

In a related point, the meditators in our study were self-selected, and while steps were taken to ensure practices met the Kabat-Zinn definition of mindfulness, and were breath or body focused, the group practiced techniques from a variety of traditions. As such, the group may have shown more heterogeneity than samples selected from a specific tradition only, and conclusions cannot be drawn regarding specific meditation techniques. However, the consistent neural activity in the meditation group shown in the topographical consistency test, combined with the significant differences in the meditation group suggest common changes in the group that were concurrent with enhanced WM performance. As such, including a variety of meditation techniques in the sample could also be viewed as a strength of the study – suggesting that the results generalise across techniques.

Another important limitation of the current study is that in our primary analyses, we were unable to examine the probe present and probe absent related neural activity separately due to an insufficient number of accepted epochs in each group (higher epoch numbers are recommended over low epoch numbers for ERP comparisons to ensure sufficient signal to noise ratio (Kappenman & Luck, 2010)). Additionally, the two groups differed in the number of epochs available for analysis, which may have the potential to influence results. To address these limitations, we performed confirmation analyses, firstly splitting the probe present and probe absent epochs to analyse them separately, and secondly excluding random epochs from the meditation group so that individuals from each group had a matched number of epochs for analysis (reported in the supplementary materials). These confirmation analyses showed the same results as the primary analyses, suggesting that these potential confounds were not the explanation for the current results.

Lastly, there are some limitations on the implications that can be drawn from the study that we think are worth noting. Although the results of this study indicate that on average the meditation group showed differences in neural activity related to WM probe presentation and decisions as well as improved WM performance, it should be noted that the results do not demonstrate that meditation reliably elicits these changes on an individual level. Medium effect sizes as per the current study suggest that the ‘average’ meditator showed better WM performance and more altered neural activity than ∼70% of the control group. However, averages reflect only the centre of a distribution. As such, some meditators showed less accurate performance than the average control. Similarly, the current results do not indicate meditation ubiquitously improves cognitive function (nor can the results answer the question of whether meditation only affects attention, or affects WM independently of attention, discussed further in the supplementary materials). Some of our research has shown null results in different measures of cognition (Bailey et al., 2018; J. Payne et al., 2019). The current research was also conducted on meditators with mostly over two years of experience, and a minimum of two hours per week of current practice, with the average amount of experience of over nine years and five hours per week of practice. As such, it is unlikely the current results reflect changes that could be obtained from a typical eight-week mindfulness course. Further research is required to fully characterise the effects of meditation across cognitive processes, at an individual level, across different durations of meditation practice, and to extend the research on the effect of meditation on WM related neural activity to clinical groups (which might show different changes to the healthy individuals sampled in the current study).

Finally, we feel it is important to note that the current results require replication before we can be confident they reflect true differences between meditators and non-meditators. Typically this point is taken for granted in scientific research, but we think it is worth noting explicitly given the current replication crisis in psychology (Collaboration, 2015), recent non replications in mindfulness research (Bailey et al., 2018; J. Payne et al., 2019), and the current media hype surrounding mindfulness research.

## Abbreviations

WM: Working Memory
EEG: Electroencephalography
MBSR: Mindfulness Based Stress Reduction
MBCT: Mindfulness Based Cognitive Therapy
FMI: Freiburg Mindfulness Inventory
FFMQ: Five Facet Mindfulness Questionnaire
ACC: Anterior Cingulate Cortex
DLPFC: Dorsolateral Prefrontal Cortex
ADHD: Attention-Deficit Hyperactivity Disorder
BAI: Beck Anxiety Inventory
BDI-II: Beck Depression Inventory II
ICA: Independent Component Analysis
AMICA: Adaptive Mixture Independent Component Analysis
RAGU: Randomisation Graphical User Interface
TCT: Topographical Consistency Test
TANOVA: Topographical Analysis of Variance
GFP: Global Field Potential
RMS: Root Mean Squared
FFT: Fast Fourier Transform
FN400: Frontal Negativity occurring ∼400 ms after probe stimuli in a working memory task
Parietal old/new effect: Positive activity in parietal regions occurring ∼600 ms after probe stimuli in a working memory task
P3: Midline parietal positivity occurring 300 to 600 ms after stimuli in many different tasks

## Supplementary Materials

The research demonstrating how the 1/f distribution could be separated from oscillatory activity was published after the current study had completed data collection, which is why hypotheses using this analysis method were exploratory. We also chose to leave examination of the WM set presentation period for future research for three reasons: Firstly, the modified Sternberg task we used contained a horizontal sequence of eight simultaneously presented letters for the WM set presentation period. Multiple simultaneously presented stimuli are not well suited for ERP analyses, and the horizontal sequence engaged horizontal eye movements which may confound neural activity comparisons. Secondly, the literature on WM activity during the WM set presentation period is sparser and less consistent, and thirdly, we had an intention to minimise multiple comparisons.

### Data Pre-processing

Second order Butterworth filtering was applied to the data with a bandpass from 1-80 Hz and a band stop filter from 47-53 Hz (data below 1 Hz was filtered out as it adversely affects the independent component analysis step). Correctly responded to trials were re-coded, and data were from the onset of the fixation cross to the offset of the visual mask for each correct trial. Single electrodes containing artifacts in more than 3% of the trials were rejected (indicated by variations in voltage larger than 250µv or kurtosis values > 5). Epochs containing artifacts were also rejected (indicated by kurtosis values > 3 for all electrodes or > 5 for single electrodes). Artifact rejections were manually verified by visual inspection by a trained EEG researcher (LRP) who was blind to group membership during the artifact rejection step. Participants were excluded if fewer than 20 correct and noise free epochs were available for analysis. This number was selected as a compromise between acceptable signal to noise ratio for analysis and maximising statistical power for the group comparisons. Research suggests cognition related ERPs and oscillations can obtain good signal to noise ratio with a minimum of 15 to 30 epochs and fewer have been used to measure oscillatory activity (Aftanas, Varlamov, Pavlov, Makhnev, & Reva, 2002; Friedman & Johnson Jr, 2000). With a minimum of 30 epochs for inclusion, another 12 participants would have been excluded.

Adaptive Mixture Independent Component Analysis (AMICA) was used to manually select and remove components related to eye movements and remaining muscle activity artifacts (Palmer, Makeig, Kreutz-Delgado, & Rao, 2008). Following the rejection of artifactual ICA components, raw data were re-filtered from 0.5-80 Hz, and all previous channel, epoch, and component rejections were re-applied, to avoid the potential effects of high pass filtering above 0.5 Hz on ERP data. Rejected electrodes were re-constructed using spherical interpolation (Perrin, Pernier, Bertrand, & Echallier, 1989). Data were then visually inspected again by a separate researcher (NWB, who was blind to the group of the data inspected at that time) to ensure the artefact rejection process was successful. Recordings were re-referenced offline to an averaged reference.

### Source Analysis

Estimation of cortical sources during topographical between-group differences was performed using Brainstorm (Tadel, Baillet, Mosher, Pantazis, & Leahy, 2011) (http://neuroimage.usc.edu/brainstorm/). EEG data were co-registered with the template model (ICBM 152) because individual MRIs were not available. The forward model used the Symmetric Boundary Element Method implemented in OpenMEEG software (Gramfort, Papadopoulo, Olivi, & Clerc, 2010). The inverse model used the computation of minimum norm estimation, with sLORETA to normalise activity based on the depth of sources (Pascual-Marqui, 2002), with dipole orientations unconstrained to the cortex to minimise the impact of using the MRI template (Lin et al., 2006). Differences in estimation were calculated using absolute subtraction. Statistical comparisons of source localisations were not performed, as scalp comparisons already demonstrated significant differences, and without MRI co-registration source statistical comparisons can be unreliable (Michel et al., 2004). Source localisation of the well-known P100 occipital ERP (averaged across 50 to 150 ms) to the correct location demonstrated our source analysis was reliable even in the absence of individual MRI templates (see Figure S5) (Malinowski, Moore, Mead, & Gruber, 2017).

**Figure S5.**
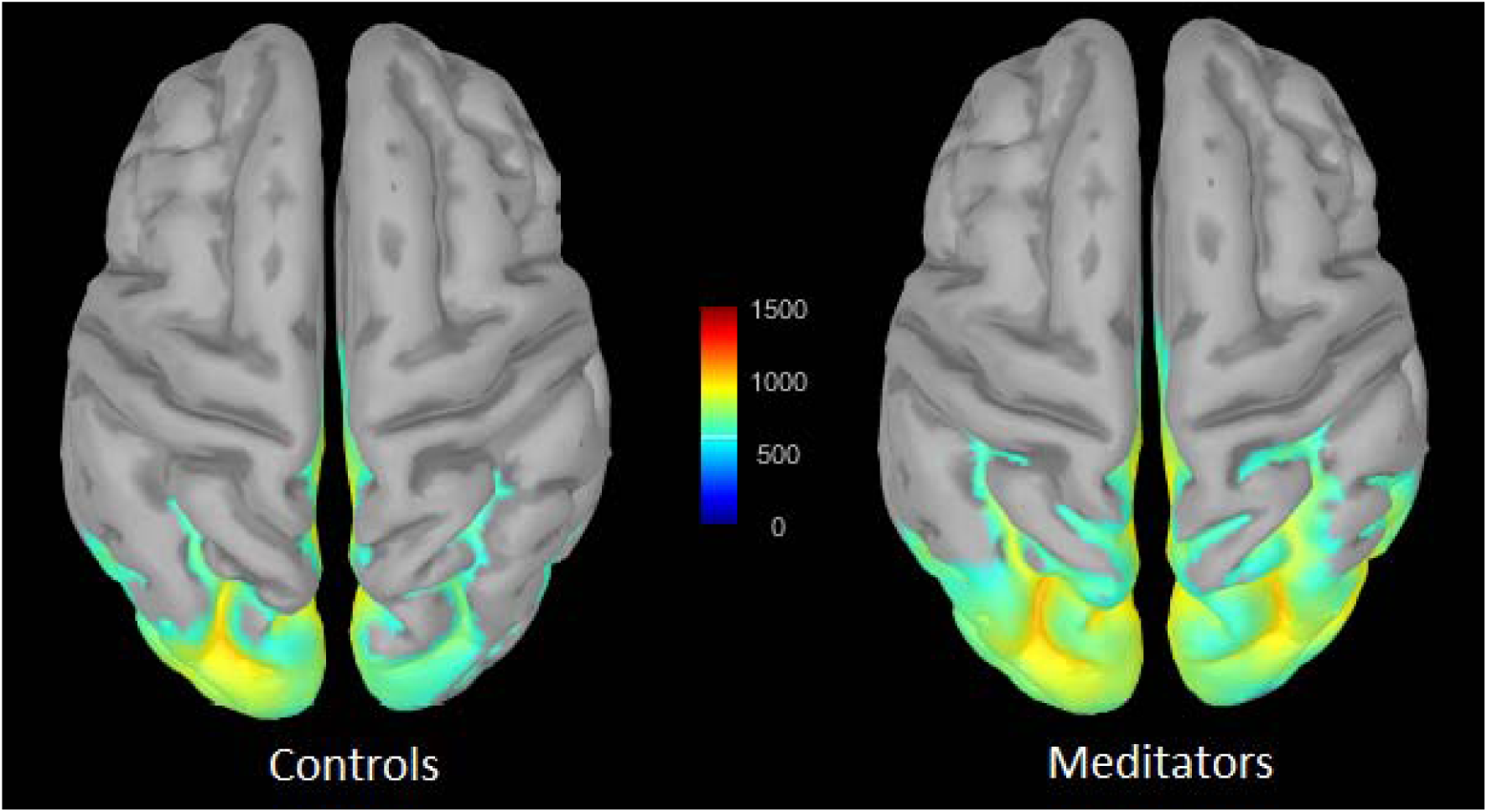
Source localised the well-known P100 occipital ERP (averaged across 50 to 150 ms) to the correct location to demonstrate our source analysis was reliable even in the absence of individual MRI templates (Malinowski et al., 2017).

### Statistical Comparisons Using RAGU

Differences between groups in overall neural response strength (across all electrodes) were compared using the GFP test, which ranks the difference between groups/conditions in GFP amplitude in the real data against the 5000 randomisations. If the real data showed larger differences than 95% of the randomised data, the results are significant at p < 0.05. Differences between groups in the distribution of activity across the scalp were compared independently of amplitude using the topographical analysis of variance (TANOVA). This test calculated the average voltage at each electrode for each group, then subtracted the values in one group from the other group to obtain a difference map. RAGU then computed the GFP of this difference map, and ranked the size of this difference map GFP from the real data against 5000 permutations of randomised data (Koenig et al., 2011). The Topographical Analysis of Covariance (TANCOVA) performed the same operations as the TANOVA, except comparisons were made between neural data and a linear predictor instead of between group comparisons (Koenig et al., 2011). Multiple comparisons were controlled for in space by collapsing values from all electrodes into a single GFP value for comparison and multiple comparisons were controlled for in time by using global duration statistics (where the duration of significant effects in the experimental data was ranked against the duration of significant effects in the randomised data). Only if the real data showed a significant effect that lasted longer than 95% of the randomised data did it pass the duration multiple comparison control. Additionally, to control for number of comparisons across the entire epoch, global count statistic p-values were measured as the sum of all significant time points in the real data across the epoch in each comparison ranked against the sum of all significant time points in the randomised data.

### Assessing 1/f Aperiodic Activity and Oscillations Separately

To assess oscillatory activity and 1/f aperiodic activity independently, 1 to 45 Hz power was calculated for each epoch across the entire delay period (5800 to 8800 ms after the start of the trial) using a Morlet Wavelet time-frequency convolution transform with 0.2 Hz resolution across sliding time windows corresponding to 5 oscillation cycles in length, with the resulting data averaged across epochs and over time in each epoch to create a measure of the power spectrum. This 1 to 45 Hz power spectrum was submitted to the Fitting Oscillations and One-Over-F (FOOOF) algorithm to extract values for frequency peaks in the theta (4 to 8 Hz) and alpha range (8 to 13 Hz), centre frequencies and bandwidths separately from 1/f slopes and offsets (Haller et al., 2018).

### Significant period in TCT test

The TCT showed significant signal indicating consistency of neural activity within all groups across the entire epoch except during the period prior to the stimulus to 80 ms after probe stimulus presentation (see Figure S1). Consistent neural activity within groups indicates that GFP and TANOVA comparisons between groups are valid.

**Figure S1.**
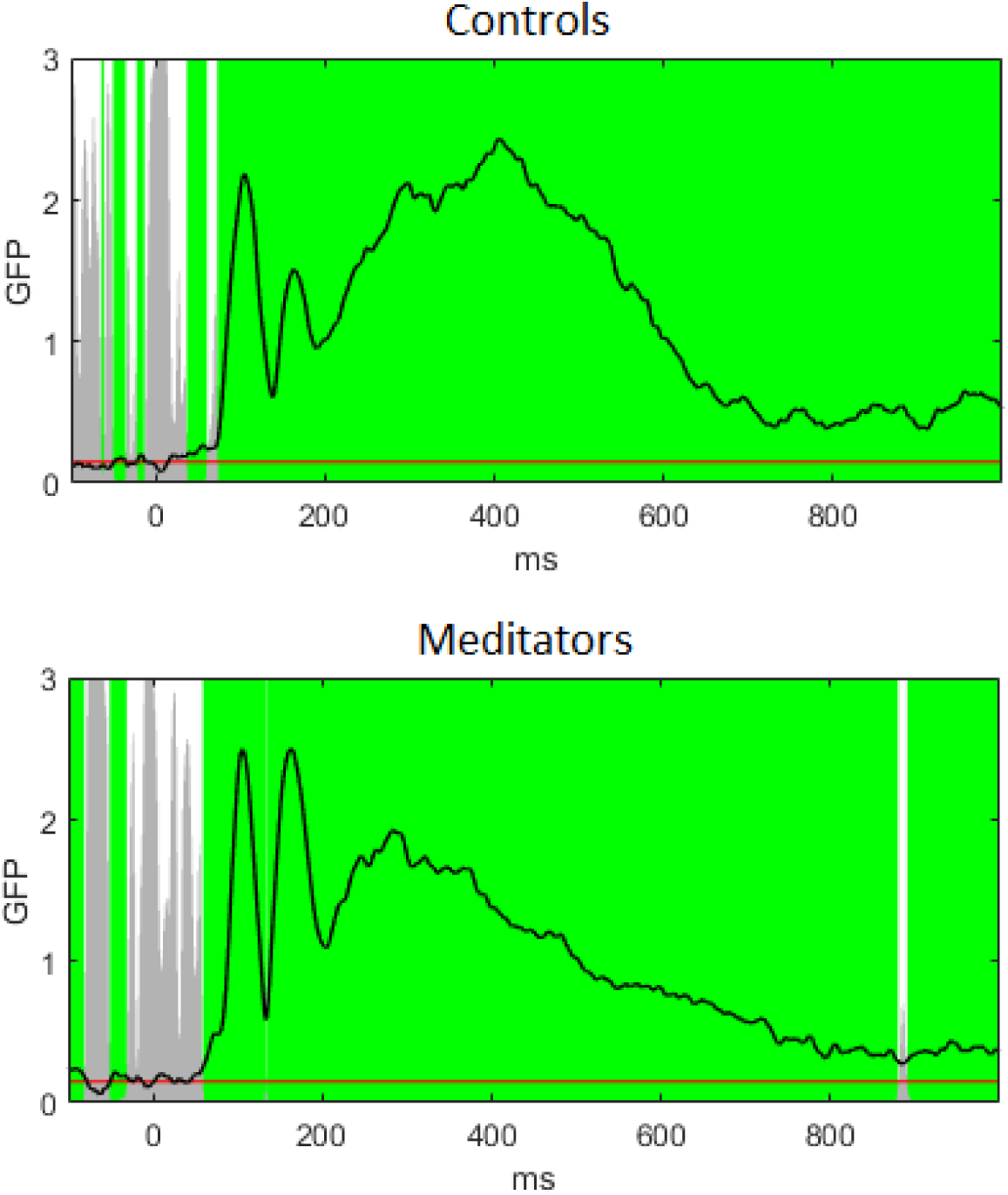
Topographical consistency test for each group. The line indicates GFP values and the grey bars indicate p-values, with the red line indicating p = 0.05. White sections indicate regions without significantly consistent distribution of activity within the group/condition, while green periods indicate consistent distribution of activity across the group/condition after duration control for multiple comparisons across time. Note significant consistency for both groups except for prior to stimulus onset to around 80 ms post stimulus presentation.

### GFP validation analyses

The number of epochs for analysis from each participant was matched between the groups by first ranking participants from each group by the number of included epochs, then randomly selecting epochs to exclude from each meditation participant until the number of epochs was matched to the control with the same epoch number rank. When only including a randomly matched number of epochs from each participant for each condition the period of significant difference in GFP between groups was reduced to 396 to 439 ms (duration control = 43 ms, p averaged across significant time period = 0.0161, η^2^ = 0.1072, see Figure S1). However, when activity was averaged across the same time period of significance as the main analysis that included all epochs (from 399 to 585 ms), the analysis restricted to a matched number of epochs from each group remained significant (p = 0.0218, η^2^= 0.0904). Additionally, splitting the conditions (and only including a matched number of epochs in each condition) provided a similar group main effect difference as the main comparisons, with a significant difference lasting from 403 to 458 ms and from 492 to 584 ms (duration control = 53ms, p-value averaged across windows of significance 403 to 458 ms = 0.0171, and p-value averaged across windows of significance 492 to 584 ms = 0.0236). No interaction between group and probe present/absent was detected (all p > 0.10). There was also a main effect of probe present/absent which was significant from 498 to 673 ms with the probe absent trials showing lower GFP amplitudes (duration control = 29ms, p-value averaged across window of significance = 0.009, η^2^ = 0.2338, η_p_^2^ = 0.2339).

**Figure S1.**
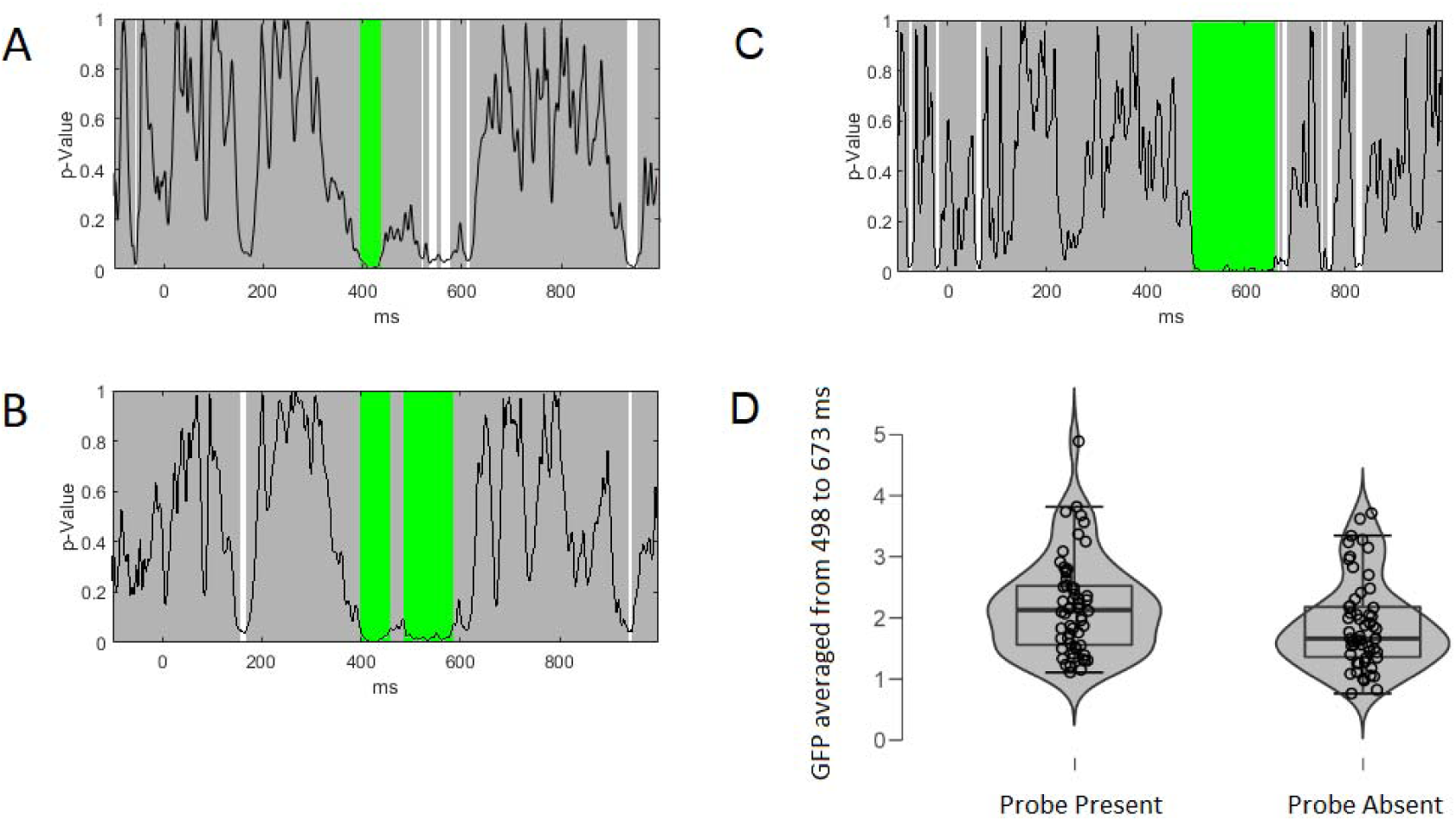
Figure S1. Validation checks for the GFP analyses after including a matched number of epochs between groups only, and splitting the conditions into probe present/absent. A – p-values across the epoch for between group comparisons including only a matched number of epochs from each group. B – p-values across the epoch for the main effect of group after splitting the epochs into probe present/absent and including this within participant factor in the comparisons. C – p-values across the epoch for the main effect of probe present/absent within participant comparisons. D – Box and violin plot of the averaged GFP across the entire window of significant difference between probe present and probe absent trials (from 498 to 673 ms). Circles reflect scores from each individual and the outer curve reflects the distribution of scores in each group (p-value averaged across window of significance = 0.009, η^2^ = 0.2338, η_p_^2^ = 0.2339).

### TANOVA validation analysis

When only including a randomly matched number of epochs from each participant for each condition the period of significant difference in TANOVA between groups remained, lasting from 290 to 628 ms (duration control = 38 ms, p averaged across significant time period = 0.0018, see Figure S2). Similarly, splitting the conditions (and only including a matched number of epochs in each condition) provided a similar group main effect difference as the main comparisons, with a significant difference lasting from 292 to 669 ms (duration control = 44 ms, p-value averaged across windows of significance = 0.0006). No interaction between group and probe present/absent was detected (all p > 0.10). There was also a main effect of probe present/absent which was significant from 333 to 478 ms and 540 to 632 ms (duration control = 26ms, p value averaged across the 333 to 478 ms window of significance = 0.0002, η^2^ = 0.0856, η_p_^2^ = 0.0862, p value averaged across the 540 to 632 ms window of significance = 0.004, η^2^ = 0.0679, η_p_^2^ = 0.0687). TCT test of the split epochs showed topographical consistency within each condition for each group (see Figure S3).

**Figure S2.**
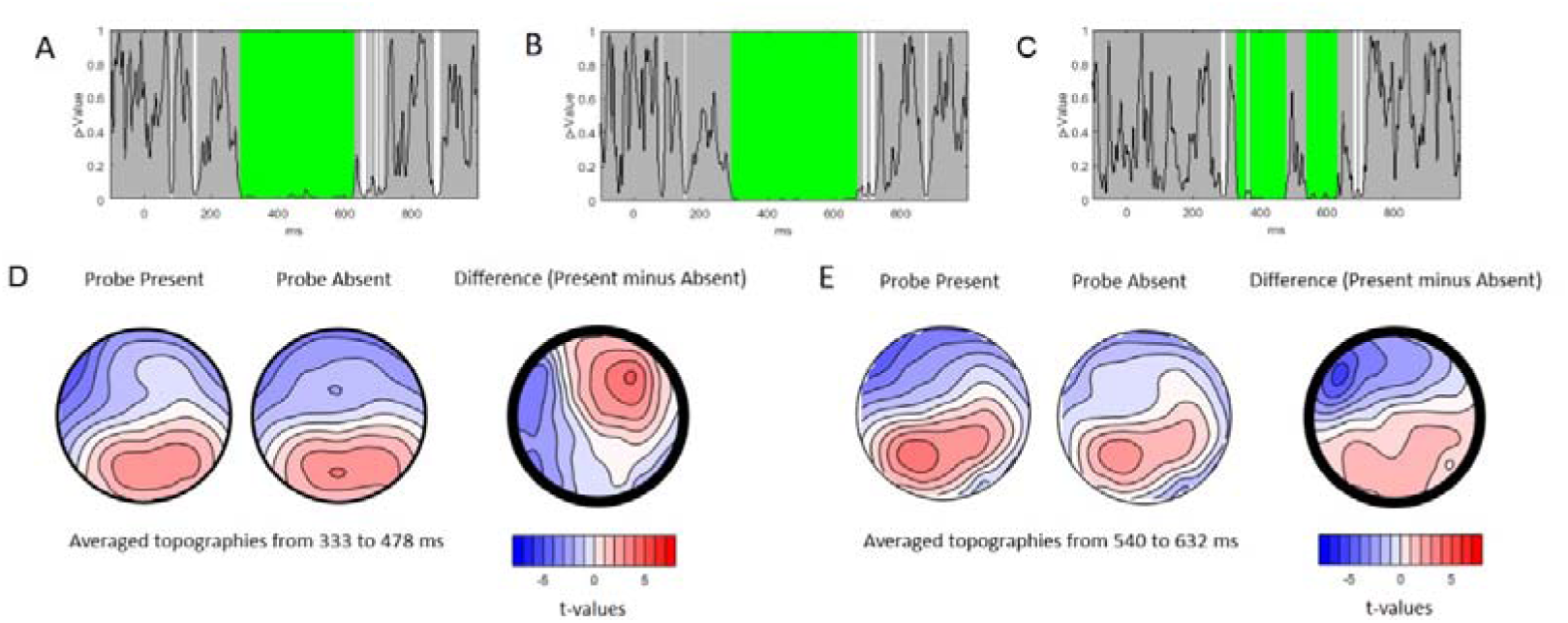
Validation checks for the TANOVA analyses after including a matched number of epochs between groups only and splitting the conditions into probe present/absent. A – p-values across the epoch for between group comparisons including only a matched number of epochs from each group. B – p-values across the epoch for the main effect of group after splitting the epochs into probe present/absent and including this within participant factor in the comparisons. C – p-values across the epoch for the main effect of probe present/absent within participant comparisons. D – Topography and difference t-maps for probe present topography minus probe absent topography during the significant time window from 333 to 478 ms (p-value. E - Topography and difference t-averaged across 333 to 478 ms window = 0.0002, η^2^ = 0.0856, η_p_^2^ = 0.0862) maps for probe present topography minus probe absent topography during the significant time window from 540 to 632 ms (p-value averaged across 540 to 632 ms window = 0.004, η^2^ = 0.0679, η_p_^2^ = 0.0687).

**Figure S3.**
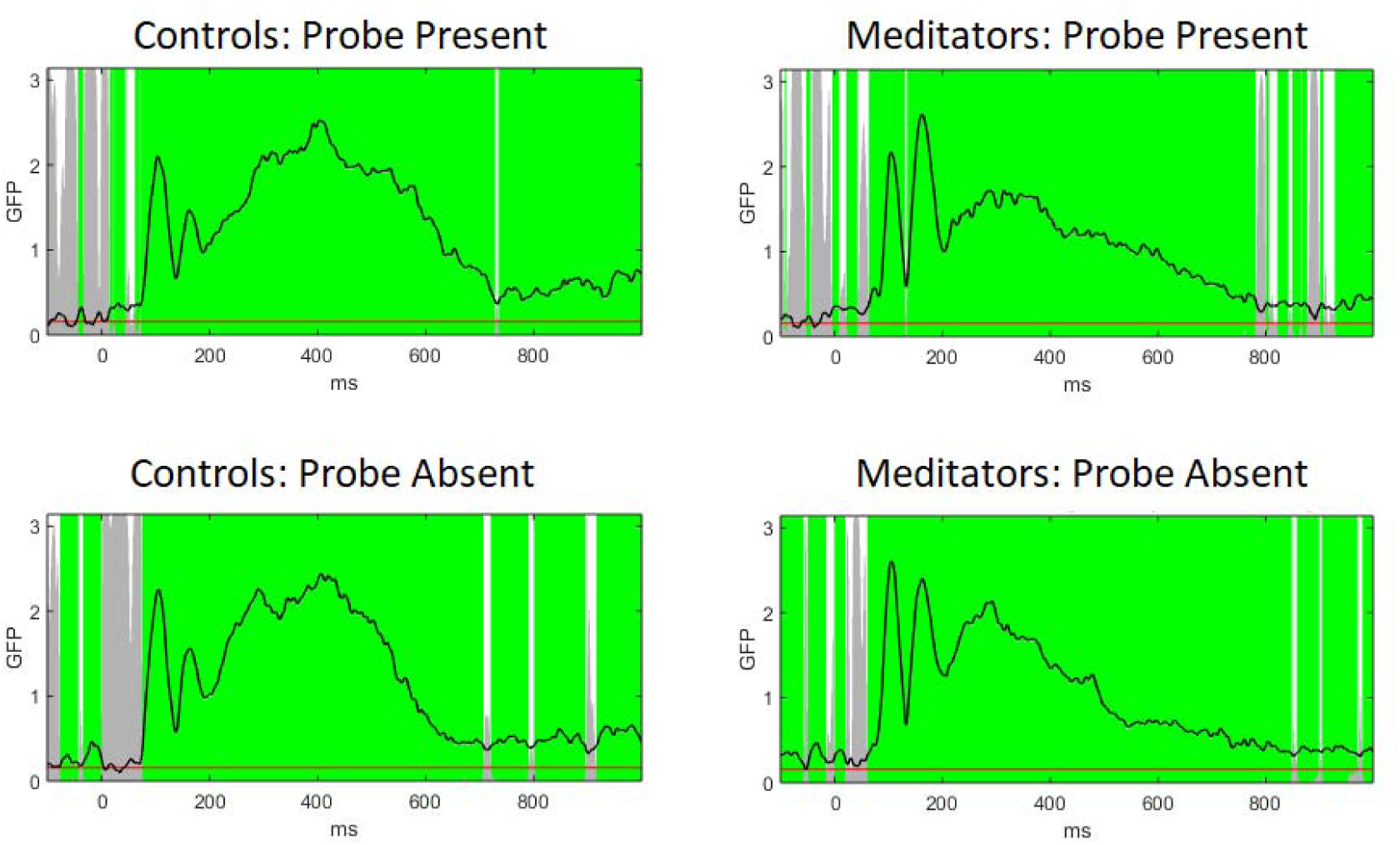
Topographical consistency test for each group and each probe present/absent condition. The line indicates GFP values and the grey bars indicate p-values, with the red line indicating p = 0.05. White sections indicate regions without significantly consistent distribution of activity within the group/condition, while green periods indicate consistent distribution of activity across the group/condition after duration control for multiple comparisons across time. Note significant consistency for both conditions in each group from 75 to 700 ms post stimulus presentation.

### Microstate Methodology

Microstates are temporarily stable topographies of neural activation that last approximately 80-120 ms before transitioning to another topography (over the course of approximately 5 ms). Different microstates reflect different source activations (Lehmann, Ozaki, & Pal, 1987). RAGU was used to identify microstates, determine the optimal number of microstates, and conduct statistical analysis of the microstates (Habermann, Weusmann, Stein, & Koenig, 2018). Microstates were identified using the atomise and agglomerate hierarchical clustering (AAHC) algorithm to merge ERP topographies into clusters, so the average topography of the clusters explains maximal variance in the ERP (Murray, Brunet, & Michel, 2008). Optimal microstate number was computed using cross-validation with the mean ERP from a learning set (N = 29) containing varied numbers of microstate classes (3-12) and timings, which are applied to the test set (N = 29) comprised of the remaining data. Smooth labelling was used for microstates, with a window size of 3 and a non-smoothness penalty of 0.5. Optimal microstate number (10) was chosen as the point where the mean variance explained in the test set reaches a plateau (Habermann et al., 2018). Randomisation statistics were then used to compare microstate properties during periods that were significant in the TANOVA.

### Exploratory Microstate Analysis of the WM Probe Period

To characterise the differences in WM probe period ERPs, we used a microstate analysis approach which clusters different time periods into dominant scalp topographies (see Figure S4). Microstate statistics were restricted to durations showing significant effects in the TANOVA. Six of the ten microstates showed differences between the groups within the significant periods from the TANOVA. Overall, the results suggest that the meditation group’s microstate map showed more right lateralized posterior positive voltages and left temporal negative voltages at around 300 ms after probe onset (in contrast to the control group’s microstate map which showed more central posterior positive voltages). After this, the meditation group’s microstate map shifted towards more frontal right positive voltages at around 400 ms after probe onset (while the control group’s microstate map showed the same shift, but not until 600 ms after probe onset).

**Figure S4.**
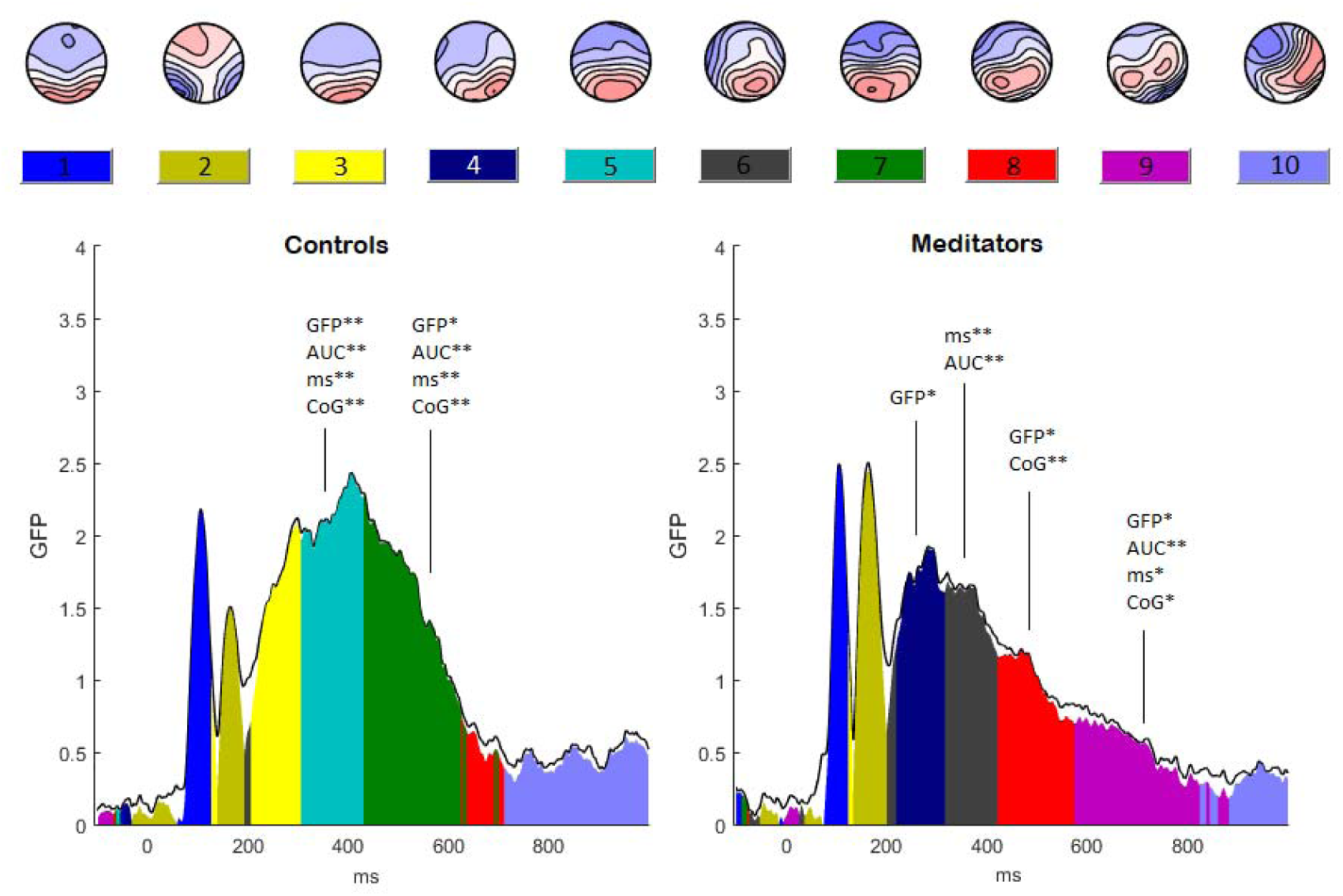
Microstate analysis between each group. GFP = mean GFP, AUC = area under the curve, ms = duration, CoG = centre of gravity, * p < 0.05, ** p = 0.01.

Microstates 4, 5 and 6 showed the distribution and time period of the FN400 (suggested to reflect familiarity and semantic processing), with meditators showing a more right posterior positive and left frontal negative FN400. Microstate 4 was only apparent in the meditation group (showing more right lateralised posterior positive voltages than microstate 3 which controls showed at the same timepoint). Microstate 4 showed a between group difference for mean GFP (p = 0.037). Microstate 6 was also only apparent in the meditation group and showed more right lateralised frontal positivity than microstate 5, which controls showed in the same time window. Microstate 6 showed differences for duration (p = 0.001) and area under the curve (p = 0.006). Microstate 5 was only apparent in the control group and showed significant differences for duration (p = 0.009), area under the curve (p = 0.006), centre of gravity (p < 0.001) and mean GFP (p < 0.001).

Microstate 7, 8 and 9 showed a distribution and time period similar to the parietal old/new effect in previous research, which has been suggested to reflect recollection. Microstate 8 showed an earlier centre of gravity (p = 0.006) and larger mean GFP (p = 0.047) in the meditator group. This microstate 8 showed more right frontal positive voltage than microstate 7’s posterior central positive voltages. Microstate 7 only appeared in the control group, and was present during the same time period as microstate 8 appeared in the meditation group. Microstate 7 showed significant differences for duration (p = 0.006), area under the curve (p = 0.001), centre of gravity (p = 0.001), and for mean GFP (p = 0.012). Microstate 9 showed even more right frontal positive voltages, and was only apparent late in the epoch in the meditation group, showing differences for duration (p = 0.015), area under the curve (p = 0.004), centre of gravity (p = 0.017), and mean GFP (p = 0.011). These differences should be considered exploratory characterisations of the differences shown during the significant periods in the TANOVA and GFP ERP analyses. While they are not controlled for multiple comparisons, they offer more detailed information about the time course and pattern of differences between the two groups in distribution of neural activity across the scalp.

### TANCOVA for Significant Differences During the WM Probe Period

Relationships between d-prime and neural activity during time windows that showed between group differences were analysed with TANCOVA. Analyses were conducted including participants in both groups to maximise statistical power and avoid increasing multiple comparisons. Analyses were performed separately for time windows reflecting distinct microstates (reported in the supplementary materials), as it was assumed that the relationship between neural activity and WM performance would be likely to vary across the three neural components that occurred during the 310 to 672 ms window differentiating the groups (microstate analyses reported above). TANCOVA comparisons showed no relationship between d-prime scores and the distribution of activity averaged across the significant time period from 310 to 425 ms (p = 0.0672), no significant relationship with the averaged activity across the significant time period from 425 to 600 ms (p = 0.1092), and no significant relationship with the significant time period from 600 to 672 ms (p = 0.1732).

### Source analysis period justification and validation of source analysis accuracy

Regarding time periods for separate depiction of source analyses, controls showed microstate 5 from 305 to 430, meditators show microstate 6 from 316 to 421 ms. We averaged the onset and offset for each to establish 310 to 425 ms for the first source analysis window. Controls showed microstate 7 from 430 to 626 ms, meditators show microstate 8 from 421 to 574 ms. We averaged the onset and offset for each to reach 425 to 600 ms. The final period for source analysis was selected based on the end of the previous period to the end of the period showing significant differences between groups in the TANOVA (600 to 672 ms).

## Supplementary theoretical points

### WM and Meditation

WM has been defined as the ability to encode, retain and manipulate a limited amount of information over the short term, an ability that is reliant on attention to bias WM processes towards goal relevant stimuli (Baddeley, 2012; Christophel et al., 2017; Ma et al., 2014; Wolpaw, 2002). WM tasks can be separated into multiple periods based on the demands required by each. Specifically, the period of exposure to the “to be remembered” stimuli can be termed “WM set presentation period”, which is separated from the period where the success of WM processes are tested (the “probe presentation period”) by the “delay period” (during which the information can be maintained via subjectively experienced strategies such as rehearsal). One additional period that could be defined is the “goal establishment period”, where participants are given instructions as to how to complete the task. The terms “WM set presentation period”, “delay period” and “probe period” are used because evidence that the brain “stores” the information during the encoding and retention periods then “retrieves” that information later has been difficult to obtain, and as such labels that suggest storage processes are misleading (Wolpaw, 2002). Rather, previous research has indicated that these periods may be associated with patterns of neural activity.

WM improvement in meditators has taken the form of increased accuracy, capacity, or prolonged periods of accurate responses in some studies and faster reaction times in others (Jha et al., 2010; Mrazek et al., 2013; Quach et al., 2016; Van Vugt & Jha, 2011; Zeidan et al., 2010). The effect has been shown in both adults and adolescents (Mrazek et al., 2013; Quach et al., 2016; Van Vugt & Jha, 2011; Zeidan et al., 2010). The effect on working memory capacity also exceeds the gains in performance obtained from increasing motivation to perform well with incentives (C. G. Jensen, Vangkilde, Frokjaer, & Hasselbalch, 2012).

Most research into WM differences in meditators has focused on how the enhanced attention from mindfulness could underpin the enhanced WM. The increased attention resulting from meditation has been suggested to increase information quality obtained in the encoding phase (Van Vugt & Jha, 2011). A modelling approach suggested this occurred via reduced perceptual noise thus increased ability in meditators to encode finer details of complex task stimuli (highly confusable faces as stimuli in a modified Sternberg task), enabling more efficient matching to or discrimination of encoded stimuli to/from the later probe stimuli during the recall period) (Van Vugt & Jha, 2011). Attentional processes have also been shown to influence neural activity in the retention periods of WM (in participants without mindfulness training), with stronger neural activity to distractor stimuli appearing in the same visual location as the WM stimuli than towards distracting stimuli appearing in other regions (Jha, 2002). Extending from this, research has shown that eight weeks of meditation training reduces visual evoked neural responses to distracting flickering of stimuli in a sustained attention and working memory task, concurrent with improved accuracy compared to the relaxation group (Schöne et al., 2018). The researchers suggested this reflected improved neural efficiency – engaging neural processes in task relevant WM information and keeping neural processes disengaged from task irrelevant information. It is also plausible that attentional processes could enhance recall, via an increased ability to attend to retrieval of the representative model of the retained encoded information (and corresponding enhancement of neural processes related to that retrieval) (Zeidan et al., 2010). A related explanation suggests that mindfulness increases meta-awareness, which mitigates the effect of mind wandering, and has been suggested to result in an increase in the number of consecutive correct responses in meditators (Zeidan et al., 2010). Indeed, research has shown that in participants who exhibit high levels of mind wandering at baseline, a reduction in mind wandering mediates the relationship between mindfulness training and increased WM capacity (Mrazek et al., 2013). This explanation is essentially an increase in the length of sustained attention, which Mrazek et al (2013) suggest is likely to be concurrent with a decrease in default mode network activation (a network associated with mind wandering).

However, it is not clear whether the positive effect of meditation on WM is simply due to attentional improvements that underpin WM, or whether mindfulness meditation has a specific effect on WM independently of the effects on attention. Buttle (2013) has argued that WM is a vital component of meditation practice, and as such WM itself is trained rather than simply attention (a perspective which has been neglected by theoretical perspectives of the mechanism of action of mindfulness). This perspective suggests that WM enables the meditator to recall the intention of the practice (focusing attention on maintaining non-judgemental awareness of sensations) (Buttle, 2011). As such, researchers have suggested that meditation could affect WM independently of attentional performance improvements, and that the WM improvements may have independent contributions as a mechanism of improved mental health resulting from meditation practice (Buttle, 2011; Jha et al., 2010). Other research has also shown meditators not to differ on measures of attention, while they did show enhanced WM performance in both short-term free recall, short term cued recall, and longer term free recall (Lykins, Baer, & Gottlob, 2012). They suggest that the lack of attentional effects may be due to good matching between groups in both age and education, and previous attention differences may have been due to poorer demographic matching, suggesting that the WM differences may reflect the true difference between meditators and controls. Some research has also suggested that while some cognitive improvements (including some, but not all, measures of attentional function such as vigilance, selective attention, and orienting) match or are smaller than those obtained by incentivising participants (suggesting the performance may be due to increased motivation rather than increased mindfulness), WM performance improvements exceeded those obtained in the non-mindful but incentivised group, and were related to changes in self-reported mindfulness scores (measured with the MAAS) (C. G. Jensen et al., 2012). Additionally, the relationship between reductions in negative affect and increased practice time has been found to be mediated by WM improvements, suggesting that WM improvements may function as a mechanism of action of meditation (rather than only attention improvements) (Jha et al., 2010).

Understanding whether meditation related attention improvement underlies WM improvements, or whether WM improvements occur above and beyond attentional improvements is important in deepening our understanding of how meditation might work. In particular, whether the primary effect of meditation is to improve attentional function, and other cognitive domains are simply influenced by that improvement, or whether the other domains might be improved independently of improvements in attentional function. The work might also provide a mechanistic justification for the application of meditation in treating mild cognitive impairment, or other conditions that show WM impairments. Exploration of neural activity related to WM may be a useful first step towards discerning between the two explanations of WM improvements in meditators. Additionally, if the results indicate strong differences between meditators and controls, this work could be further developed for use as a biomarker of improved neural function as a result of meditation, and potentially contribute to a mechanism of action understanding of how the practice of mindfulness meditation can lead to improved clinical outcomes – it has been suggested that WM capacity benefits emotional regulation (Schmeichel, Volokhov, & Demaree, 2008) and higher WM ability in meditators has been associated with reduced negative emotion (Jha et al., 2010).

### Discussion supplementary points

Source analysis indicated reduced activity in fronto-midline, central midline and left central regions, in the parietal old/new effect ERP, and reduced left temporal activity in the later part of the component, as well as increased activity in right occipital regions. These reductions in the later part of the ERP likely reflect a combination of the TANOVA and GFP results, as the source analysis does not normalise for amplitude when comparing distributions (unlike the TANOVA).

Although our results indicated no differences in the WM delay period, it may be that other neural processes in either the WM stimuli presentation period or WM delay period differ in meditators to enable the enhanced performance. Indeed, it seems strange that the only differences in neural activity would be in the WM probe period, particularly when meditation has been suggested to increase information quality obtained in the encoding phase (Van Vugt & Jha, 2011). One possible example worth exploring has been demonstrated by Tran, Hoffner, LaHue, Tseng, and Voytek (2016), who found that although older adults showed a reduced ability to modulate alpha activity during WM delay periods in ipsilateral visual regions to WM stimuli concurrent with decreased WM performance compared to younger adults, this activity was unrelated to the WM performance decreases, which were only predicted by phase synchronisation of visual region alpha phase to the onset of alerting stimuli. Research into meditators has indicated they show more synchronisation of the phase of neural activity to stimuli or to cues to anticipate stimuli (as suggested previous research (Lutz et al., 2009; Slagter, Lutz, Greischar, Nieuwenhuis, & Davidson, 2009), however also see (J. Payne et al., 2019)). These or similar attention related changes to the anticipation of WM stimuli, to the WM stimuli presentation period, or to the WM delay period may enable the enhanced WM performance without differences in alpha or theta power during the WM delay period. Additionally, it may be that oscillatory changes underlie or are enabled by the WM probe period ERP changes found in the current study. Previous research has suggested sub-delta activity modulation underpins WM related P3 activity, and that P3 amplitude changes are associated with changes to induced theta amplitudes, possibly reflecting specific activation of WM relevant regions of the cortex while the P3 reflects inhibition of widespread non-specific regions (W Klimesch et al., 2000).

However, ultimately, it may be difficult to detect differences between meditators and controls in the WM delay period of a WM task because the theoretical perspective underlying the experiment is flawed. The approach taken may implicitly assume there is a neural signal reflecting the storage of information, something that has been argued not to exist and to reflect a misunderstanding of how the brain connects the past and present – through synaptic activity, neurotransmitter concentrations, dendritic branching and membrane properties rather than “storing” information (Wolpaw, 2002). Perhaps we have detected no differences in the WM delay period because we are looking for storage that is not there, and research would do better to focus on the process linking previous experience to current behaviour (Wolpaw, 2002).

Additionally, it is worth noting that different WM tasks and different stimuli are likely to elicit other differences between meditators and controls, as there is good evidence for stimulus selectivity in WM based on sensory specific activation, and activity detected in this study does not reflect simply storage of WM information in the form of letters, but also timing information to help with anticipation, stimulus-response pairing, mnemonic strategies, and other processes, all of which generate different associated neural activity (Christophel et al., 2017; Riley & Constantinidis, 2016).

### Enhanced WM processes vs Enhanced Attention processes?

The increased left temporal lobe activity in meditators does appear at face value to indicate altered WM specific function. This area is not traditionally associated with attention, and is traditionally associated with WM. However, P3 activity has been associated with attention (which is generated by temporal regions as well as other regions and occurs during the same time window as the temporal lobe differences in the meditation group). Additionally, even if meditators showed altered activity in a WM specific area that is not related to attention, there is no way to determine that that activity is not the downstream result of earlier attention related differences in neural activity. Our view is that to answer the question of whether meditation specifically effects WM without attention modulations, we need a better definition of attention as performed by the brain (in contrast to the psychological model of attention). Our suspicion is that when we have a brain-based definition of attention function, separating WM and attention functions may become meaningless. This is because our developing view of how the brain performs the function that we have labelled “attention” is that it modulates neural activity as best it can in whatever manner is necessary to meet task demands. For example, variation in attentional ability might be reflected by increased ability to generate fronto-midline theta to resolve conflicts between different neural processes in a Stroop or Go Nogo task, or modulate alpha activity to inhibit distracting information being processed in a non-task-relevant brain region in a task with a tactile distractor (Wang et al., 2019), or in the current study, earlier temporal activation in the recall period. While research suggests that attention is related to frontal and parietal activity to maintain focus, we suspect activity in these regions reflects only part of the story and may not even be active in all attention modulations. Instead, we suggest “that the differences in meditators reflect improved attentional function, and this improved attentional function provides enhancements to neural processes that are the ‘weakest link’ in achieving task-oriented goals (Lavie, 1995; Rauss, Pourtois, Vuilleumier, & Schwartz, 2009; Vogel, Woodman, & Luck, 2005). Attention supports the processes most likely to fail in the chain from stimulus processing to response, reducing the chance of failure at those most vulnerable points and enhancing the probability of successful task performance.” (N. W. Bailey et al., 2019).

## References

Aftanas, L. I., Varlamov, A. A., Pavlov, S. V., Makhnev, V. P., & Reva, N. V. (2002). Time-dependent cortical asymmetries induced by emotional arousal: EEG analysis of event-related synchronization and desynchronization in individually defined frequency bands. International Journal of Psychophysiology, 44(1), 67–82.

Baddeley, A. (2012). Working memory: theories, models, and controversies. Annual review of psychology, 63, 1–29.

Baer, R. A., Smith, G. T., Hopkins, J., Krietemeyer, J., & Toney, L. (2006). Using self-report assessment methods to explore facets of mindfulness. Assessment, 13(1), 27–45.

Bailey, N., Freedman, G., Raj, K., Sullivan, C., Rogasch, N., Chung, S., … Van Dam, N. (2019). Mindfulness meditators show altered distributions of early and late neural activity markers of attention in a response inhibition task. PLOS ONE, 14(8), e0203096.

Bailey, N. W., Freedman, G., Raj, K., Sullivan, C. M., Rogasch, N. C., Chung, S. W., … Van Dam, N. T. (2019). Mindfulness meditators show altered distributions of early and late neural activity markers of attention in a response inhibition task. PLOS ONE, 14(8), e0203096.

Bailey, N. W., Raj, K., Freedman, G., Fitzgibbon, B. M., Rogasch, N. C., Van Dam, N. T., & Fitzgerald, P. B. (2018). Mindfulness meditators do not show differences in electrophysiological measures of error processing. Mindfulness, 1–21.

Beck, A. T., Steer, R. A., & Brown, G. K. (1996). Beck depression inventory-II. San Antonio, 78(2), 490–498.

Benjamini, Y., & Hochberg, Y. (1995). Controlling the false discovery rate: a practical and powerful approach to multiple testing. Journal of the Royal statistical society: series B (Methodological), 57(1), 289–300.

Britton, W. B., Davis, J. H., Loucks, E. B., Peterson, B., Cullen, B. H., Reuter, L., … Lindahl, J. R. (2018). Dismantling Mindfulness-Based Cognitive Therapy: Creation and validation of 8-week focused attention and open monitoring interventions within a 3-armed randomized controlled trial. Behaviour research and therapy, 101, 92–107.

Brookes, M. J., Wood, J. R., Stevenson, C. M., Zumer, J. M., White, T. P., Liddle, P. F., & Morris, P. G. (2011). Changes in brain network activity during working memory tasks: a magnetoencephalography study. Neuroimage, 55(4), 1804–1815.

Buttle, H. (2011). Attention and Working Memory in Mindfulness–Meditation Practices. The Journal of Mind and Behavior, 123–134.

Buttle, H. (2013). More than the sum of my parts: a cognitive psychologist reflects on mindfulness/meditation experience. Reflective Practice, 14(6), 766–773.

Cavanagh, J. F., & Frank, M. J. (2014). Frontal theta as a mechanism for cognitive control. Trends in cognitive sciences, 18(8), 414–421.

Chang, Y. K., Huang, C. J., Chen, K. F., & Hung, T. M. (2013). Physical activity and working memory in healthy older adults: an ERP study. Psychophysiology, 50(11), 1174–1182.

Christophel, T. B., Klink, P. C., Spitzer, B., Roelfsema, P. R., & Haynes, J.-D. (2017). The distributed nature of working memory. Trends in cognitive sciences, 21(2), 111–124.

Collaboration, O. S. (2015). Estimating the reproducibility of psychological science. Science, 349(6251), aac4716.

Curran, T., & Cleary, A. M. (2003). Using ERPs to dissociate recollection from familiarity in picture recognition. Cognitive Brain Research, 15(2), 191–205.

Delorme, A., & Makeig, S. (2004). EEGLAB: an open source toolbox for analysis of single-trial EEG dynamics including independent component analysis. Journal of neuroscience methods, 134(1), 9–21.

Duarte, A., Ranganath, C., Winward, L., Hayward, D., & Knight, R. T. (2004). Dissociable neural correlates for familiarity and recollection during the encoding and retrieval of pictures. Cognitive Brain Research, 18(3), 255–272.

Edwards, J., Peres, J., Monti, D. A., & Newberg, A. B. (2012). The neurobiological correlates of meditation and mindfulness. In Exploring frontiers of the mind-brain relationship (pp. 97–112): Springer.

Ergen, M., Yildirim, E., Uslu, A., Gürvit, H., & Demiralp, T. (2012). P3 response during short-term memory retrieval revisited by a spatio-temporal analysis. International Journal of Psychophysiology, 84(2), 205–210.

Federmeier, K. D., & Laszlo, S. (2009). Time for meaning: Electrophysiology provides insights into the dynamics of representation and processing in semantic memory. Psychology of learning and motivation, 51, 1–44.

Finnigan, S., Humphreys, M. S., Dennis, S., & Geffen, G. (2002). ERP ‘old/new’effects: memory strength and decisional factor (s). Neuropsychologia, 40(13), 2288–2304.

Friedman, D., & Johnson Jr, R. (2000). Event-related potential (ERP) studies of memory encoding and retrieval: A selective review. Microscopy research and technique, 51(1), 6–28.

Gao, R., Peterson, E. J., & Voytek, B. (2017). Inferring synaptic excitation/inhibition balance from field potentials. Neuroimage, 158, 70–78.

Gramfort, A., Papadopoulo, T., Olivi, E., & Clerc, M. (2010). OpenMEEG: opensource software for quasistatic bioelectromagnetics. Biomedical engineering online, 9(1), 45.

Gu, B.-M., van Rijn, H., & Meck, W. H. (2015). Oscillatory multiplexing of neural population codes for interval timing and working memory. Neuroscience & Biobehavioral Reviews, 48, 160–185.

Guglietti, C. L., Daskalakis, Z. J., Radhu, N., Fitzgerald, P. B., & Ritvo, P. (2013). Meditation-related increases in GABAB modulated cortical inhibition. Brain stimulation, 6(3), 397–402.

Habermann, M., Weusmann, D., Stein, M., & Koenig, T. (2018). A Student’s Guide to Randomization Statistics for Multichannel Event-Related Potentials Using Ragu. Frontiers in neuroscience, 12.

Haller, M., Donoghue, T., Peterson, E., Varma, P., Sebastian, P., Gao, R., … Voytek, B. (2018). Parameterizing neural power spectra. bioRxiv, 299859.

Hayama, H. R., Johnson, J. D., & Rugg, M. D. (2008). The relationship between the right frontal old/new ERP effect and post-retrieval monitoring: specific or non-specific? Neuropsychologia, 46(5), 1211–1223.

Hutzler, F. (2014). Reverse inference is not a fallacy per se: Cognitive processes can be inferred from functional imaging data. Neuroimage, 84, 1061–1069.

Jensen, C. G., Vangkilde, S., Frokjaer, V., & Hasselbalch, S. G. (2012). Mindfulness training affects attention—or is it attentional effort? Journal of Experimental Psychology: General, 141(1), 106.

Jensen, O., & Tesche, C. D. (2002). Frontal theta activity in humans increases with memory load in a working memory task. European Journal of Neuroscience, 15(8), 1395–1399.

Jha, A. P. (2002). Tracking the time-course of attentional involvement in spatial working memory: An event-related potential investigation. Cognitive Brain Research, 15(1), 61–69.

Jha, A. P., Stanley, E. A., Kiyonaga, A., Wong, L., & Gelfand, L. (2010). Examining the protective effects of mindfulness training on working memory capacity and affective experience. Emotion, 10(1), 54.

Kabat-Zinn, J. (2009). Wherever you go, there you are: Mindfulness meditation in everyday life: Hachette Books.

Kappenman, E. S., & Luck, S. J. (2010). The effects of electrode impedance on data quality and statistical significance in ERP recordings. Psychophysiology, 47(5), 888–904.

Kerr, C. E., Sacchet, M. D., Lazar, S. W., Moore, C. I., & Jones, S. R. (2013). Mindfulness starts with the body: somatosensory attention and top-down modulation of cortical alpha rhythms in mindfulness meditation. Frontiers in human neuroscience, 7, 12.

Klimesch, W., Doppelmayr, M., Schwaiger, J., Winkler, T., & Gruber, W. (2000). Theta oscillations and the ERP old/new effect: independent phenomena? Clinical Neurophysiology, 111(5), 781–793.

Klimesch, W., Sauseng, P., & Hanslmayr, S. (2007). EEG alpha oscillations: the inhibition–timing hypothesis. Brain research reviews, 53(1), 63–88.

Koenig, T., Kottlow, M., Stein, M., & Melie-García, L. (2011). Ragu: a free tool for the analysis of EEG and MEG event-related scalp field data using global randomization statistics. Computational Intelligence and Neuroscience, 2011, 4.

Kottlow, M., Schlaepfer, A., Baenninger, A., Michels, L., Brandeis, D., & Koenig, T. (2015). Pre-stimulus BOLD-network activation modulates EEG spectral activity during working memory retention. Frontiers in behavioral neuroscience, 9, 111.

Lagopoulos, J., Xu, J., Rasmussen, I., Vik, A., Malhi, G. S., Eliassen, C. F., … Holen, A. (2009). Increased theta and alpha EEG activity during nondirective meditation. The Journal of Alternative and Complementary Medicine, 15(11), 1187–1192.

Lavie, N. (1995). Perceptual load as a necessary condition for selective attention. Journal of Experimental Psychology: Human perception and performance, 21(3), 451.

Lehmann, D., Ozaki, H., & Pal, I. (1987). EEG alpha map series: brain micro-states by space-oriented adaptive segmentation. Electroencephalography and clinical neurophysiology, 67(3), 271–288.

Lin, F.-H., Witzel, T., Ahlfors, S. P., Stufflebeam, S. M., Belliveau, J. W., & Hämäläinen, M. S. (2006). Assessing and improving the spatial accuracy in MEG source localization by depth-weighted minimum-norm estimates. Neuroimage, 31(1), 160–171.

Lutz, A., Slagter, H. A., Dunne, J. D., & Davidson, R. J. (2008). Attention regulation and monitoring in meditation. Trends in cognitive sciences, 12(4), 163–169.

Lutz, A., Slagter, H. A., Rawlings, N. B., Francis, A. D., Greischar, L. L., & Davidson, R. J. (2009). Mental training enhances attentional stability: neural and behavioral evidence. Journal of Neuroscience, 29(42), 13418–13427.

Lykins, E. L., Baer, R. A., & Gottlob, L. R. (2012). Performance-based tests of attention and memory in long-term mindfulness meditators and demographically matched nonmeditators. Cognitive therapy and research, 36(1), 103–114.

Ma, W. J., Husain, M., & Bays, P. M. (2014). Changing concepts of working memory. Nature neuroscience, 17(3), 347.

MacLean, K. A., Ferrer, E., Aichele, S. R., Bridwell, D. A., Zanesco, A. P., Jacobs, T. L., … Shaver, P. R. (2010). Intensive meditation training improves perceptual discrimination and sustained attention. Psychological science, 21(6), 829–839.

Malinowski, P., Moore, A. W., Mead, B. R., & Gruber, T. (2017). Mindful aging: the effects of regular brief mindfulness practice on electrophysiological markers of cognitive and affective processing in older adults. Mindfulness, 8(1), 78–94.

Manning, J. R., Jacobs, J., Fried, I., & Kahana, M. J. (2009). Broadband shifts in local field potential power spectra are correlated with single-neuron spiking in humans. Journal of Neuroscience, 29(43), 13613–13620.

Maurer, U., Brem, S., Liechti, M., Maurizio, S., Michels, L., & Brandeis, D. (2015). Frontal midline theta reflects individual task performance in a working memory task. Brain topography, 28(1), 127–134.

Michel, C. M., Murray, M. M., Lantz, G., Gonzalez, S., Spinelli, L., & de Peralta, R. G. (2004). EEG source imaging. Clinical Neurophysiology, 115(10), 2195–2222.

Miller, K. J., Hermes, D., Honey, C. J., Hebb, A. O., Ramsey, N. F., Knight, R. T., … Fetz, E. E. (2012). Human motor cortical activity is selectively phase-entrained on underlying rhythms. PLoS computational biology, 8(9), e1002655.

Mrazek, M. D., Franklin, M. S., Phillips, D. T., Baird, B., & Schooler, J. W. (2013). Mindfulness training improves working memory capacity and GRE performance while reducing mind wandering. Psychological science, 24(5), 776–781.

Murray, M. M., Brunet, D., & Michel, C. M. (2008). Topographic ERP analyses: a step-by-step tutorial review. Brain topography, 20(4), 249–264.

Nagel, B. J., Herting, M. M., Maxwell, E. C., Bruno, R., & Fair, D. (2013). Hemispheric lateralization of verbal and spatial working memory during adolescence. Brain and cognition, 82(1), 58–68.

Osaka, M., Komori, M., Morishita, M., & Osaka, N. (2007). Neural bases of focusing attention in working memory: an fMRI study based on group differences. Cognitive, Affective, & Behavioral Neuroscience, 7(2), 130–139.

Palmer, J. A., Makeig, S., Kreutz-Delgado, K., & Rao, B. D. (2008). Newton method for the ICA mixture model. Paper presented at the 2008 IEEE International Conference on Acoustics, Speech and Signal Processing.

Pascual-Marqui, R. D. (2002). Standardized low-resolution brain electromagnetic tomography (sLORETA): technical details. Methods Find Exp Clin Pharmacol, 24(Suppl D), 5–12.

Payne, J., Baell, O., Geddes, H., Fitzgibbon, B., Emonson, M., Hill, A., … Bailey, N. (2019). Experienced meditators exhibit no differences to demographically-matched controls in theta phase synchronisation, P200, or P300 during an auditory oddball task. bioRxiv, 608547.

Payne, L., & Kounios, J. (2009). Coherent oscillatory networks supporting short-term memory retention. Brain research, 1247, 126–132.

Perrin, F., Pernier, J., Bertrand, O., & Echallier, J. (1989). Spherical splines for scalp potential and current density mapping. Electroencephalography and clinical neurophysiology, 72(2), 184–187.

Peterson, E. J., Rosen, B. Q., Campbell, A. M., Belger, A., & Voytek, B. (2018). 1/f neural noise is a better predictor of schizophrenia than neural oscillations. bioRxiv, 113449.

Peterson, E. J., & Voytek, B. (2017). Alpha oscillations control cortical gain by modulating excitatory-inhibitory background activity. bioRxiv, 185074.

Poldrack, R. A. (2006). Can cognitive processes be inferred from neuroimaging data? Trends in cognitive sciences, 10(2), 59–63.

Pontifex, M. B., Hillman, C. H., & Polich, J. (2009). Age, physical fitness, and attention: P3a and P3b. Psychophysiology, 46(2), 379–387.

Quach, D., Mano, K. E. J., & Alexander, K. (2016). A randomized controlled trial examining the effect of mindfulness meditation on working memory capacity in adolescents. Journal of Adolescent Health, 58(5), 489–496.

Rac-Lubashevsky, R., & Kessler, Y. (2018). Oscillatory correlates of control over working memory gating and updating: An eeg study using the reference-back paradigm. Journal of cognitive neuroscience, 30(12), 1870–1882.

Rauss, K. S., Pourtois, G., Vuilleumier, P., & Schwartz, S. (2009). Attentional load modifies early activity in human primary visual cortex. Human brain mapping, 30(5), 1723–1733.

Riley, M. R., & Constantinidis, C. (2016). Role of prefrontal persistent activity in working memory. Frontiers in systems neuroscience, 9, 181.

Rugg, M. D., & Curran, T. (2007). Event-related potentials and recognition memory. Trends in cognitive sciences, 11(6), 251–257.

Rugg, M. D., & Henson, R. N. (2002). Episodic memory retrieval: an (event-related) functional neuroimaging perspective. In: Psychology Press.

Rugg, M. D., Mark, R. E., Walla, P., Schloerscheidt, A. M., Birch, C. S., & Allan, K. (1998). Dissociation of the neural correlates of implicit and explicit memory. Nature, 392(6676), 595.

Sauseng, P., Griesmayr, B., Freunberger, R., & Klimesch, W. (2010). Control mechanisms in working memory: a possible function of EEG theta oscillations. Neuroscience & Biobehavioral Reviews, 34(7), 1015–1022.

Sauseng, P., Hoppe, J., Klimesch, W., Gerloff, C., & Hummel, F. C. (2007). Dissociation of sustained attention from central executive functions: local activity and interregional connectivity in the theta range. European Journal of Neuroscience, 25(2), 587–593.

Scheeringa, R., Petersson, K. M., Oostenveld, R., Norris, D. G., Hagoort, P., & Bastiaansen, M. C. (2009). Trial-by-trial coupling between EEG and BOLD identifies networks related to alpha and theta EEG power increases during working memory maintenance. Neuroimage, 44(3), 1224–1238.

Schmeichel, B. J., Volokhov, R. N., & Demaree, H. A. (2008). Working memory capacity and the self-regulation of emotional expression and experience. Journal of personality and social psychology, 95(6), 1526.

Schöne, B., Gruber, T., Graetz, S., Bernhof, M., & Malinowski, P. (2018). Mindful breath awareness meditation facilitates efficiency gains in brain networks: A steady-state visually evoked potentials study. Scientific reports, 8(1), 13687.

Sheehan, D. V., Lecrubier, Y., Sheehan, K. H., Amorim, P., Janavs, J., Weiller, E., … Dunbar, G. C. (1998). The Mini-International Neuropsychiatric Interview (MINI): the development and validation of a structured diagnostic psychiatric interview for DSM-IV and ICD-10. The Journal of clinical psychiatry.

Slagter, H. A., Lutz, A., Greischar, L. L., Nieuwenhuis, S., & Davidson, R. J. (2009). Theta phase synchrony and conscious target perception: impact of intensive mental training. Journal of cognitive neuroscience, 21(8), 1536–1549.

Steer, R., & Beck, A. (1997). Beck anxiety inventory.

Sternberg, S. (1966). High-speed scanning in human memory. Science, 153(3736), 652–654.

Stróżak, P., Abedzadeh, D., & Curran, T. (2016). Separating the FN400 and N400 potentials across recognition memory experiments. Brain research, 1635, 41–60.

Tadel, F., Baillet, S., Mosher, J. C., Pantazis, D., & Leahy, R. M. (2011). Brainstorm: a user-friendly application for MEG/EEG analysis. Computational Intelligence and Neuroscience, 2011, 8.

Tallon-Baudry, C., Bertrand, O., Delpuech, C., & Pernier, J. (1997). Oscillatory γ-band (30–70 Hz) activity induced by a visual search task in humans. Journal of Neuroscience, 17(2), 722–734.

Tang, Y.-Y., Ma, Y., Fan, Y., Feng, H., Wang, J., Feng, S., … Li, J. (2009). Central and autonomic nervous system interaction is altered by short-term meditation. Proceedings of the National Academy of Sciences, 106(22), 8865–8870.

Tang, Y.-Y., Ma, Y., Wang, J., Fan, Y., Feng, S., Lu, Q., … Fan, M. (2007). Short-term meditation training improves attention and self-regulation. Proceedings of the National Academy of Sciences, 104(43), 17152–17156.

Tran, T. T., Hoffner, N. C., LaHue, S. C., Tseng, L., & Voytek, B. (2016). Alpha phase dynamics predict age-related visual working memory decline. Neuroimage, 143, 196–203.

Van Vugt, M. K., & Jha, A. P. (2011). Investigating the impact of mindfulness meditation training on working memory: A mathematical modeling approach. Cognitive, Affective, & Behavioral Neuroscience, 11(3), 344–353.

Vigneau, M., Beaucousin, V., Herve, P.-Y., Duffau, H., Crivello, F., Houde, O., … Tzourio-Mazoyer, N. (2006). Meta-analyzing left hemisphere language areas: phonology, semantics, and sentence processing. Neuroimage, 30(4), 1414–1432.

Vogel, E. K., Woodman, G. F., & Luck, S. J. (2005). Pushing around the locus of selection: Evidence for the flexible-selection hypothesis. Journal of cognitive neuroscience, 17(12), 1907–1922.

Voytek, B., Kramer, M. A., Case, J., Lepage, K. Q., Tempesta, Z. R., Knight, R. T., & Gazzaley, A. (2015). Age-related changes in 1/f neural electrophysiological noise. Journal of Neuroscience, 35(38), 13257–13265.

Wagner, A. D., Shannon, B. J., Kahn, I., & Buckner, R. L. (2005). Parietal lobe contributions to episodic memory retrieval. Trends in cognitive sciences, 9(9), 445–453.

Walach, H., Buchheld, N., Buttenmüller, V., Kleinknecht, N., & Schmidt, S. (2006). Measuring mindfulness—the Freiburg mindfulness inventory (FMI). Personality and individual differences, 40(8), 1543–1555.

Wang, M. Y., Freedman, G., Raj, K., Fitzgibbon, B. M., Sullivan, C., Tan, W.-L., … Bailey, N. W. (2019). Mindfulness meditation alters neural activity underpinning working memory during tactile distraction. bioRxiv, 790584.

Winawer, J., Kay, K. N., Foster, B. L., Rauschecker, A. M., Parvizi, J., & Wandell, B. A. (2013). Asynchronous broadband signals are the principal source of the BOLD response in human visual cortex. Current Biology, 23(13), 1145–1153.

Wolpaw, J. R. (2002). Memory in neuroscience: rhetoric versus reality. Behavioral and cognitive neuroscience reviews, 1(2), 130–163.

Woodruff, C. C., Hayama, H. R., & Rugg, M. D. (2006). Electrophysiological dissociation of the neural correlates of recollection and familiarity. Brain research, 1100(1), 125–135.

Yonelinas, A. P. (2002). The nature of recollection and familiarity: A review of 30 years of research. Journal of memory and language, 46(3), 441–517.

Zeidan, F., Johnson, S. K., Diamond, B. J., David, Z., & Goolkasian, P. (2010). Mindfulness meditation improves cognition: Evidence of brief mental training. Consciousness and cognition, 19(2), 597–605.

